# A theory of subicular function and generalized vector coding

**DOI:** 10.64898/2026.04.04.716474

**Authors:** Fei Wang, Andrej Bicanski

**Affiliations:** Max-Planck Institute for Human Cognitive and Brain Sciences, Department of Psychology, Leipzig, Germany

**Keywords:** Subiculum, Spatial memory, Cognitive map, Corner cells, Boundary vector cells

## Abstract

Located in the hippocampal formation, the subiculum exhibits numerous, seemingly unrelated neural codes, without a common account of neuronal selectivity. We propose Disco, a theory that can explain activity in the subiculum and related neural codes in other areas. Given the continuous flow of experience, subicular activity can be understood as signalling discontinuities in that flow, coded in a vector-base. Geometric discontinuities drive responses to walls, drops, objects, holes, corners and curved surfaces. The generality of Disco enables predictions for 3D environments, natural scenes, and mixed-modality coding. Non-spatial discontinuities in behavioral state yield axis-coding, and tracking population activity signals event boundaries as temporal discontinuities. The ability to generate this diverse set of neural activity suggests that discontinuity coding is a key principle underlying subicular activity.

## 1 Introduction

Knowledge of environmental boundaries, barriers, and objects is paramount for navigation and memory (1–3). Explicit neural codes for allocentric (Fig. 1A) and egocentric vector representations to boundaries and objects have first been reported in rodents (4–8). Anatomically, these vectorial codes are localized primarily in the subiculum, commonly thought of as a region which conveys the results of hippocampal processing to other brain areas (9–13). Human data corroborates the rodent findings regarding boundaries (14–16). The rodent discoveries followed on the heels of theoretical predictions, both for boundaries (17) and objects (18). Interestingly, the subiculum also exhibits so-called corner cells (3) that fire selectively when the animal is near any corner\object in the environment (Fig. 1B), and axis coding (19) along movement trajectories.

**Fig. 1.**
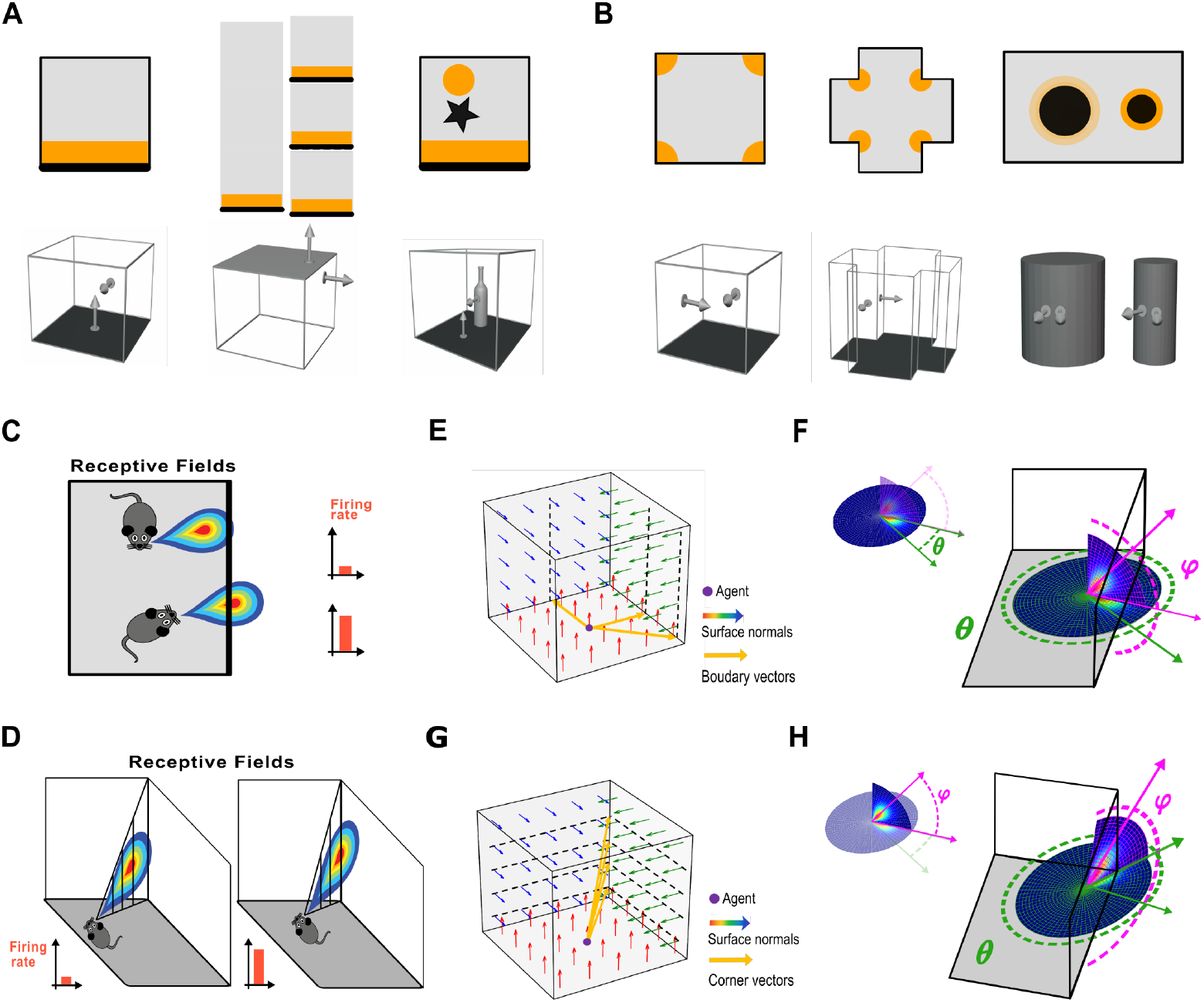
The Disco hypothesis unifies vector coding of diverse geometric features. (**A-B**) Representative subicular firing patterns. Top: geometric environments, with orange regions indicating neural firing when binned by animal location. Bottom: proxies for geometric features in terms of surface normals. Boundary vector cells (BVCs) respond to walls(5), drops(31) (formed by joining or separating three unwalled platforms), and objects(21) (A). All of these elements are characterized by abrupt changes in surface normals between the floor and their surfaces. Corner cells(3) respond to concave and convex corners as well as to objects (B). Corners are defined by abrupt changes in surface normals between two neighboring walls. Differences in firing rates between high- and low-convexity regions reflect the rate of change of surface normals on the object’s surface (B, bottom panels), with color intensity indicating firing rate magnitude. (**C**) BVC receptive fields (attached to cartoon mice) are selective to environmental boundaries at a specific distance and azimuth. (**D**) Corner cells encode corners at a specific distance and elevation relative to the animal. (**E-F**) Model schematic of BVCs: a boundary as segmentation between walls and floor (E). The tuning vector is defined by the agent’s relative azimuth (*θ*) and distance to the boundary, which is detected as the discontinuity in the elevation (*φ*) component of adjacent surface normals (F). (**G-H**) Model schematic of corner cells: a corner as segmentation between vertical walls (G). The tuning vector is defined by the agent’s relative elevation and distance, which is detected as the discontinuity in the azimuth component of adjacent surface normals (H).

However, a large gap in our understanding regarding these neural codes persists. While we can model both boundary and object vector coding with the same receptive field equations (17,18) (Fig. 1C), it remains unknown how the brain might compute these functions. What do objects and boundaries have in common to facilitate this coding scheme? This question becomes more puzzling when we appreciate that even drops (Lever et al., 2009; Stewart et al., 2014) (e.g., at the edge of a table), or a traversable stripe on the floor (which is not a navigational impediment) are recognized as “barriers” (21). This can be compared to the well-defined neural selectivity observed in the sensory periphery (e.g., orientation-selective cells in the visual cortex (22)). Prior models of boundary vector coding have been highly useful in modelling spatial cognition but stipulate specific selectivity, and the presence of a boundary/object is given to the model (5,17,18,23) (Fig. 1C). Moreover, subicular neurons coding for corners (3) and reference axes (19) appear unrelated.

Here, we introduce the coding of discontinuities of experience as a general principle for subicular function – the Disco model. We propose that all of the above cell types are essentially discontinuity detectors, signaling salient changes in space and likely other modalities (i.e., non-spatial discontinuities). That is, given the continuous flow of experience, subicular activity can be understood as signaling discontinuities in that flow. In the simplest formulation of this idea (Fig. 1A), a boundary is a discontinuity from the floor to the wall or a drop. An object is a discontinuity from the floor to the object surface. When we appreciate that animals move in 3 dimensions (often neglected for the sake of modelling, cf. Fig. 1C), corner cells can be framed as discontinuities between adjacent vertical walls (Fig. 1B). When the walls separate, the corner field disappears (3) because the specific type of discontinuity has disappeared.

We apply Disco to a series of experimental setups (fig. S1) and show that major subicular cell types (as discovered in the rodents) emerge as different discontinuities of experience, including joint boundary vector (BVCs) (5) and object vector cells (4) (OVCs), height-dependent corner cells (3) (in convex, concave, and continuous flavor). Generalizing to non-geometric domains, Disco yields axis-tuned cells (in behavioral/velocity space) and BVC responses to traversable stripes (combining geometric and color/texture space). We also show that event boundaries (24,25) can manifest as discontinuities in the population activity. Additionally, we demonstrate that Disco can be applied to situations with complex and noisy geometric features, enabling predictions of neural activity during navigation in natural environments – a first for boundary, object and corner coding, and a stark departure from the need of prior models to stipulate object/boundary presence and to explicitly define what element of an environment should be considered a driver of neural activity.

In the spatial domain, these discontinuities can be parsimoniously modelled using normal vectors of perceived surfaces (Fig. 1, E and G) (also see reference (26)). A priori it remains an open question if the brain explicitly calculates normal vectors or uses other indicators. The nature of the discontinuity signal remains an open question. However, the fact that so many cell types can emerge from the same coding principle, suggests it is not simply a convenient algorithmic proxy. Taken together, our results show that it is possible to formulate a domain-general framework that captures a wide spectrum of spatial and non-spatial cell types in the subiculum and potentially other areas. Disco is compatible with broader hippocampal function and prior theoretical accounts of hippocampal memory, suitably reframed as indexing the discontinuities of experience (27,28), the privileged anatomical position of the subiculum (different discontinuities need to be communicated to distinct cortical output targets) (9,11,29), and the notion of a cortical cascade of memory (unpacking discontinuities to specific exemplar representations) (30).

## 2 Results

### 2.1 Model overview

A full description of the model is relegated to the materials and methods. Geometric codes can illustrate the relatedness and complementarity of subicular selectivity for boundaries/objects and corners when framed as discontinuity detection in 3D. The activity of BVCs is paradigmatically described in a vector-coding base (17,18) (Fig. 1C), structured as a product of Gaussians (𝒢) for azimuth (*θ*; 0◦: East, 90◦: North, 180◦: West, 270◦: South) and distance (*l*). This model captures BVC activity when the allocentric location of the boundary can be supplied (Fig. 1C). Two crucial additions are made here to generalize subicular responses. First, extending the two-dimensional (2D) receptive fields into three dimensions. Second, Disco augments the vector-coding framework with a detector, specifying which type of discontinuity is sampled by the cell or its upstream inputs (Fig. 1, E and F). For boundaries, *b* in Equation (1) captures the process of boundary identification.

We assume the BVC is selective for discontinuities with respect to elevation (*φ*; Upward, 180◦: Downward, 0◦) at a fixed azimuth and distance. Thus, Disco provides a generalized account of various BVC-tuned elements (e.g., walls, objects, and drops), which can be signaled by discontinuities in perceived surface normals. For each location *<u>x</u>* in the environment, the firing of a BVC is given by

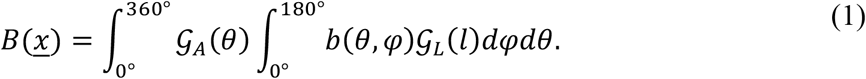

Here 𝒢_*L*_ refers to Gaussian selectivity for distance, while 𝒢_*A*_ signifies the Gaussian selectivity for azimuth. *b*(*θ, φ*) is the boundary detector driven by changes in sampled surface normals. By removing the integration over *φ* and directly supplying the locations of present boundary discontinuities (e.g., a square box, where, *b*(*θ, φ* → 90◦) = 1), the model can be reduced to the previously proposed BVC function (17). Thus, the present formulation provides a mechanistic account of the detection of boundaries and other features at the specific locations.

Swapping elevation and azimuth in Equation (1), we derive tuning functions for corner cells (Fig. 1, D to H). Here the implicit assumption is that corner cells (and in fact other subicular neurons) are neurons that sample a different combination of environmental discontinuities from upstream regions as compared to BVCs, a notion that will also be important for mixed-modality coding (see below). The corner detector (*c* in Equation (2)) represents the selectivity to azimuthal discontinuities in perceived surface normals. The vector-coding base of corner cells is a product of Gaussians over elevation and distance (Fig. 1, H).

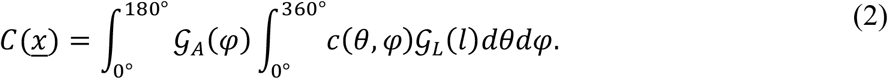

### 2.2 Boundary vector responses across varied experiments

Disco can generate BVC activities across a wide range of laboratory settings (Fig. 2, A and B). BVCs driven by the detection of discontinuities in surface normals respond to boundaries in a square, and exhibit a field doubling after barrier insertion. In a cylinder, the surface normals along the wall vary smoothly, but the discontinuity with respect to elevation at a fixed azimuth, still occurs at the wall-floor interface. Likewise, the curved surface of an object does not disturb BVC selectivity for the discontinuity from the floor to the object surface.

**Fig. 2.**
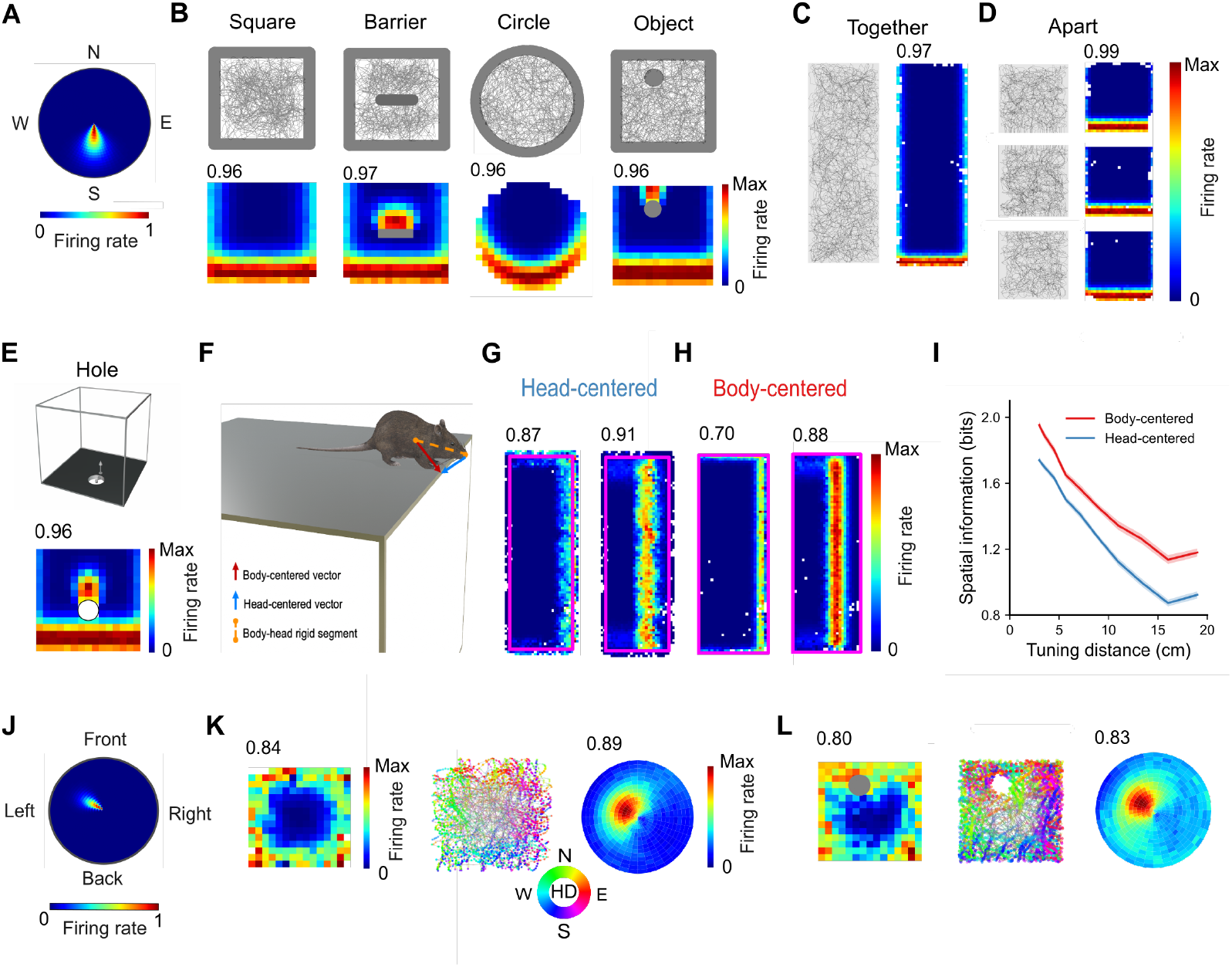
Boundary and Object vector responses emerge from the detection of discontinuities. Example BVC receptive field for a discontinuity located at a short distance south of the agent. Top: Schematics of environmental configurations and simulated trajectories. Bottom: Rate maps of example BVCs, (color-coded by firing rate) showing responses to square walls, an inserted barrier, circular walls, and an object. (**C-D**) BVC responses to drops in the simulated “together– apart” manipulation, where three unwalled square platforms (C) form a single rectangular open platform, which are then separated (D). The gaps between platforms remain traversable. The neuron responds to discontinuities detected at the platform edges. (**E**) Predicted BVC responses to a hole due to abrupt changes in surface normals at the edge. (**F**) An extended two-point agent model allowing the “head” to extend beyond the platform edge. Through the rigid body–head reference frame change, the vector from the head to a discontinuity is transformed into a body-centered vector, which determines the firing rate of BVCs. (**G-H**) Rate maps of example BVCs with short (left) and long (right) tuning distances. When computed from head position, firing fields can extend beyond the platform edge (pink line) (G). Body-centered rate maps are constrained to the platform (H), conveying more spatial information (**I**). (**J**) Receptive field of the example egocentric boundary cell (EBC) in the Disco framework. (**K-L**) EBC responses to walls and objects. Left, allocentric rate maps. Middle, trajectory plots with neuronal firing color-coded by head direction, with transparency proportional to firing rate. Right, egocentric boundary/object rate maps, where firing rate is represented as a function of proximal boundary/object locations in polar coordinates relative to the agent.

Boundaries have previously been defined by a combination of sensory cues and movement constraints (5). Compared to upright vertical surfaces, drops provide very different sensory cues, and are often considered as movement-limiting boundaries. However, BVCs also fire to the drop between two raised platforms even when rats can jump across it (no change to the transition structure, cf. Fig.4 in reference (31)). Here, we simulated BVC activity in this “together–apart’’ manipulation experiment (31), where initially joined platforms are separated. Simulated BVCs responded to drops, detecting the discontinuity in surface normals from the floor to the platform edge (Fig. 1A), reproducing drop-elicited field repetition (Fig. 2, C and D). The counterpart to an edge in the object case is a hole in the floor. Disco predicts that BVCs should show vector coding to holes in the floor (Fig. 2E). These results re-iterate that Disco provides a generalized account for the diverse aspects of neural selectivity of BVCs (21).

Contrary to walls, drops introduce the possibility that an animal can lean over the drop edge. To simulate this setting, we extended the agent model to a two-point variant: one point tracks the head and the other the body, with the body constrained to remain on the platform (materials and methods). Experimental data show that rate maps based on head position reveal two BVC subtypes, encoding vectors to (i) the platform edge and (ii) the outer limit of space reachable by the animal’s head (5). However, the standard BVC tuning function cannot account for this classification, as the tuning distance and direction become ambiguous. E.g., should the direction be north or south if the head extends beyond the northern wall? In our model, the head detects the discontinuity of the environment; a head–body coordinate transform then yields the boundary vector from the body to the discontinuity (Fig. 2F). When we draw the rate map based on the head position, BVCs with tuning distances shorter than the head–body separation can extend their fields beyond the platform edge (Fig. 2G). Thus, the two-point extension predicts that BVC vector coding is fundamentally body-centered, reconciling the experimental observation mentioned above (5). Moreover, body-based rate maps exhibit more precise positional coding (Fig. 2, G to I).

Disco principles can also be leveraged to build egocentric boundary coding (6,8,32). By constructing receptive fields in an egocentric reference frame (Fig. 2J), Disco egocentric boundary cells (EBCs) exhibited stable wall-bearing spatial tuning in a square box, responding when a wall occurred at a specific distance and azimuth relative to the agent’s head direction (Fig. 2K). Using the simulated data, we also constructed previously established egocentric boundary rate maps (6) to characterize the egocentric vector representation of boundaries when the neuron fires. Rather than using wall locations, as in previous study, we computed the relative azimuths and distances of discontinuities (materials and methods). The measured rate maps reflect the underlying receptive fields that mechanistically govern neuronal firing, but remain imperfect due to variations in exploratory trajectories (Fig. 2K). Disco EBCs also perform egocentric bearing of objects (33), showing that objects can be detected as object–floor discontinuities in the egocentric frame (Fig. 2L). The egocentric vector-base could still be subject to transformation to the allocentric frame as in prior models (18,23).

### 2.3 Discrete and continuous corner cell responses

Evolving naturally from the BVC account, we propose an explicit tuning function for subicular corner cells based on Disco framework (materials and methods), consistent with the subiculum’s comprehensive representation of spatial geometry (3). Disco detects the presence of corners by recognizing the discontinuity of surface normals at wall junctions (Fig. 1B). As the corner vectors are defined by relative elevation and distance (Fig. 1G), these cells can fire near environmental corners in all azimuthal orientations. Corner cells also fire near circular walls (3) with surface normals rotating continuously with respect to azimuth. Disco reproduces the activity observed in these continuous geometries (Fig. 3A) by capturing the discontinuities from finite-resolution sampling (materials and methods). I.e., surface normals are not sampled at infinitely close locations.

**Fig. 3.**
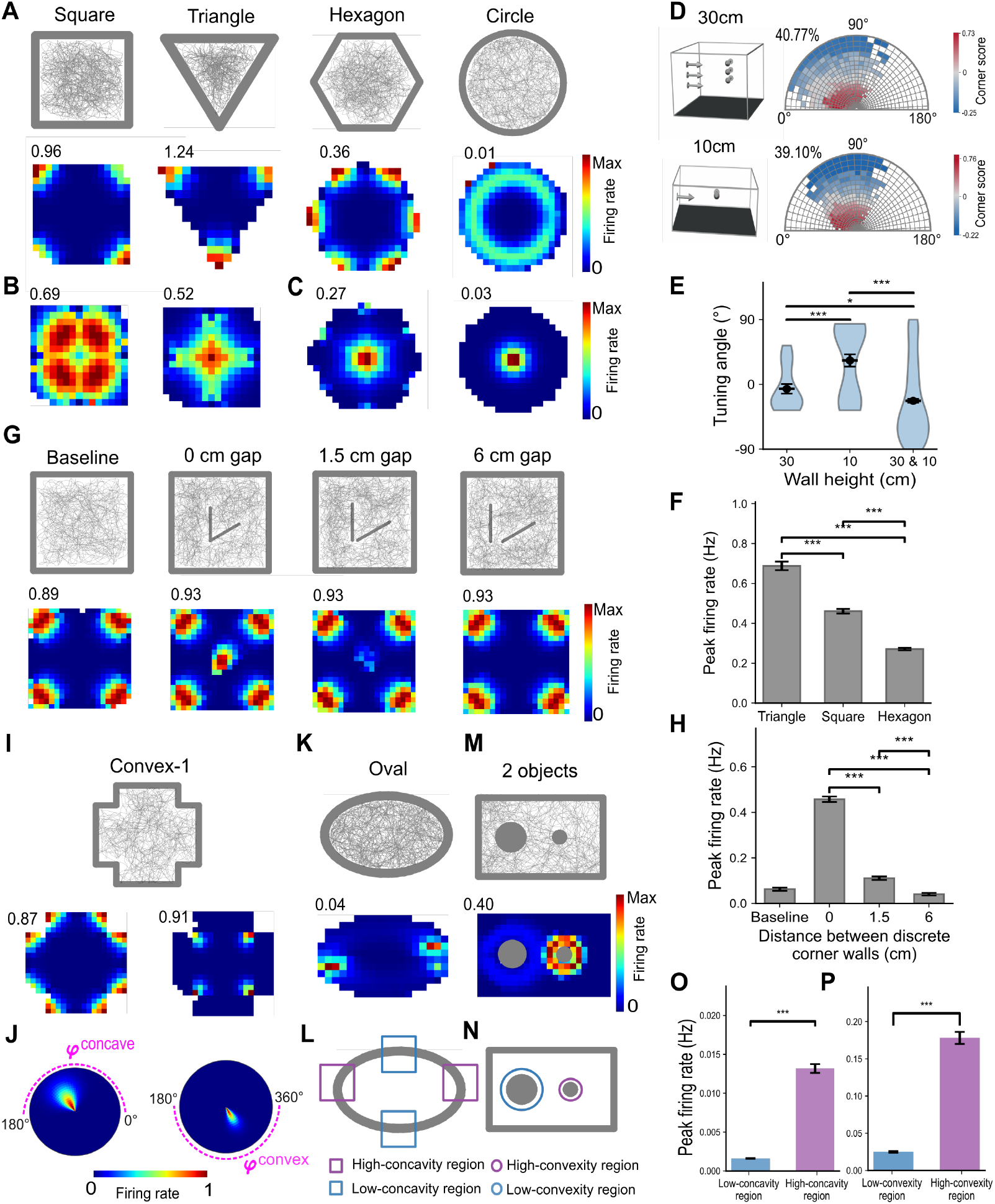
Corner cells naturally emerge from the Disco formalism. (**A**) Open-arena environment shapes and an example corner cell, plotted as in Fig. 2b. This neuron responds to a discontinuity located at a short distance (2.2 cm) and positive elevation relative to the agent. (**B-C**) Two example neurons with positive tuning elevations and long tuning distances in the open-field arena. Tuning distances are 14.6 cm for (B, left) and 24.5 cm for (B, right) and (C), respectively. (**D**) Schematic of high-wall and low-wall square arenas and the distribution of corner scores across neurons. The proportion of corner cells (corner score > 0) is indicated in the upper left. (**E**) Tuning angles of non-corner cells in square environments with different wall heights. Violin plots depict the density distribution; black dots with error bars indicate mean ± s.e.m. (30, *n* = 20 neurons; 10, *n* = 33 neurons; 30&10, *n* = 439 neurons). (**F**) Peak firing rates of corner cells for each environment (Triangle, *n* = 288 neurons ; square, *n* = 316 neurons ; hexagon, *n* = 315 neurons ; mean ± s.e.m.). (**G**) Open-field arenas with an inserted discrete corner and an example corner cell. (**H**) Peak firing rates of baseline-identified corner cells around the inserted corner across all sessions (*n* = 319 neurons ; mean ± s.e.m.). (**I**) Arenas with concave and convex corners and two representative corner cells encoding concave (bottom left) and convex (bottom right) corners. (**J**) The two corner cells in i exhibit non-overlapping receptive fields. (**K**) Oval arena and the concave corner cell (I). (**L**) Illustration showing high- and low-concavity regions. (**M**) Arena containing two cylindrical objects with convex corner cell responses (I). (**N**) Illustration showing high- and low-convexity regions. (**O-P**) Peak firing rates of active neurons in low- and high-concavity (convexity) regions (Low-concavity, *n* = 138 neurons; high-concavity, *n* = 608 neurons; low-convexity, *n* = 467 neurons; high-convexity, *n* = 567 neurons; mean ± s.e.m.) *P < 0.05, **P < 0.01, ***P < 0.001.

Similar to experimental studies, we adopted the corner score (3,34) to classify corner cells. The score ranged from −1 for fields at the arena center to 1 for fields precisely at a corner. A neuron was classified as a corner cell if its corner score was greater than 0; otherwise, it was classified as a non-corner cell (materials and methods). Neurons with positive tuning elevations and shorter tuning distances were preferentially classified as corner cells (Fig. 3D, and fig. S2. Here it can be emphasized that cells not formally classified as corner cells are still governed by the corner-cell receptive fields as defined above. They simply don’t pass the experimental criterion, and could be interpreted as representing different geometric features. In regular-shaped environments (Fig. 3A, see irregular-shape simulation in fig. S3), the firing fields of these cells progressively merged as tuning distance increased. Hence, in square arenas distal corner tuning (of nominally non-corner cells) can represent the principal spatial axes (Fig. 3B, right). Cells with intermediate distance tuning resemble the reported conjunctive boundary-vertex code in retrosplenial cortex (RSC), which was hypothesized to emerge from distally tuned vertex cells (35). Incidentally, vertex cells could also be modelled within the Disco framework (supplementary text). In arenas with more corners, corner fields converged on the geometric center (Fig. 3C), and in a circular arena became reminiscent of center-distance cells (32,36,37) whose firing rates are positively correlated with distance to the geometric centroid.

We further tested whether Disco captures the three defining features of corners: (i) wall height, (ii) corner angle, and (iii) the junction of two walls. The proportion of subiculum corner cells decreased from the high-wall (30cm) to the low-wall (10cm) square (3). Disco predicts that this is due to taller walls providing more detectable discontinuities (Fig. 3D). In the low-wall square, more neurons with positive tuning elevations were classified as non-corner cells (30cm versus 10cm: *t*_41.4_ = 1.29, *P* = 0.21, *d* = 0.36 ; 30cm versus 30&10cm: *t*_44.1_ = −2.96, *P* = 0.00, *d* = −0.43; 10cm versus 30&10cm: *t*_23.6_ = −4.03, *P* = 0.00, *d* = −0.72; 2-sample t test) (Fig. 3E), as their receptive fields likely extended above the wall height during agent’s exploration. Tuning distances of non-corner cells did not differ significantly between the two conditions (fig. S4).

Acute corner angles elicited higher firing rates (3). Disco offers a natural explanation: smaller corner angles correspond to larger differences between the two adjacent surface normals. We used their dot product to control the amplitude of firing rate (materials and methods), thereby introducing a modulation by corner angles (Triangle versus square: *t*_440.8_ = 9.31, *p* = 0.00, *d* = 0.768 ; triangle versus hexagon: *t*_332.9_ = −2.96, *P* = 0.00, *d* = 1.55 ; square versus hexagon: *t*_479.6_ = 14.59, *P* = 0.00, *d* = 1.16; 2-sample t test) (Fig. 3F).

Next, we simulated the experimental paradigm in which mice explored a square box with an inserted discrete corner (3). Consistent with BVCs’ response to the inserted barrier (Fig. 2B), Disco corner cells formed new firing fields near the inserted corner (Fig. 3G), which weakened as the two connected walls were gradually separated and the corresponding discontinuities disappeared (0 cm gap versus 1.5 cm: *t*_670.2_ = 23.69, *P* = 0.00, *d* = 1.69 ; 0cm versus 6cm: *t*_554.8_ = 30.74, *P* = 0.00, *d* = 2.20; 1.5cm versus 6cm: *t*_710.4_ = 7.12, *P* = 0.00, *d* = 0.51; 2-sample t test) (Fig. 3H). However, experiments demonstrate that corner cells still exhibited weak firing near the separated walls as animals continued to perceive them as a corner (3). Disco can resolve this ambiguity by broadening the notion of discontinuity (see supplementary text). Corner detection initially considered only jump discontinuities (materials and methods), where both limits exist but differ (e.g., the surface normals of two connected walls). By including the second type of discontinuity, where one limit does not exist (e.g., undefined normal vector “in the air”), firing at wall endpoints can be generated (fig. S5). Alternatively, the corner response after separation could be a trace phenomenon, reflecting mnemonic input from CA1 (21,38).

Experimental data (3) showed that convex and concave corners are encoded by separate neuronal populations (Fig. 3, I and J). We defined a subset of corner cells whose tuning vectors oppose those representing concave corners. Algorithmically, Disco distinguishes concave from convex corners using the cross-product direction of adjacent surface normals, enabling each subset to encode one corner type (materials and methods). Combining this cross-product classification with the dot-product measure of curvature magnitude, Disco reproduces the subiculum’s broader coding scheme for concavity and convexity, including in continuous environments (3). Concave corner cells showed higher firing rates in high-concavity regions in an oval enclosure (low-versus high-concavity: *t*_612.6_ = −20.42, p = 0.00, d = −1.17; 2-sample t test) (Fig. 3, K, L, and O). Convex corner cells display spatial coding surrounding the objects, unlike BVC’s directional tuning to objects (Fig. 1A, Fig. 2B), and exhibit higher firing rates around a highly convex cylindrical object (low-versus high-convexity: *t*_584.1_ = −18.67, p = 0.00, d = −1.11; 2-sample t test) (Fig. 3, M, N, and P).

Our formalism extends to egocentric corner coding (figs. S6, see supplementary text), which yields the tuning function of egocentric vertex cells (when relaxing the effects of concavity/convexity and corner angles) observed in the RSC (35). Unlike EBCs (Fig. 2, J to L), this coding arises from constraining discontinuity detection to a specific angular window. We also studied alternative models for corner cells based on the same input as BVCs (supplementary text). Corner selectivity can be generated by ring-like receptive fields (fig. S7) or by receiving a thresholded sum of BVC inputs (35) (figs. S8–S11). These models are capable of reproducing corner representations but do not to explain the effects of external factors (e.g., corner angles, concavity/convexity, curvature). This further supports our hypothesis regarding corner cells’ selectivity to the elevation angle (Fig. 1D and H), implying that detecting corners and objects requires the animal’s up-and-down scanning (39). Sun et al. demonstrated that cutting tactile inputs did not affect corner cell activity, whereas darkness had a significant impact (3).

### 2.4 Emergence of geometric coding in a natural environment

As shown above, Disco suggests a common class of triggers (detected discontinuities) as drivers of subicular activity. This allows Disco to go beyond typical laboratory environments composed of planes (e.g., floors and walls). Natural terrains typically exhibit non-flat topography. How terrain features such as hills, basins, and irregular surfaces shape the activity of spatial/vector cells (40–42) is unknown. The challenge lies in defining geometric features in the real world. Should a stone be considered an object/boundary or should it be ignored? Should small, irregular environmental features be represented? This ambiguity prevents prior models, in which boundary/object presence must be specified by the modeler, from being applied in three-dimensional (3D), realistic settings, limiting the ecological validity. Disco provides a workflow for modelling navigation cells in natural environments, without the need to explicitly specify geometric features.

Before applying Disco to a natural scene, we extended the model to account for more variable sensory information in 3D environments. Animal trajectories are largely confined to a horizontal plane in experiments. We first simulated an agent exploring bowl-like and sink-like environments. In the bowl, multiple discontinuities of surface normals along the same horizontal direction are detected, generating multiple boundary-vectors (Fig. 4A). This superposition can be modeled as the average of these boundary vectors (materials and methods). Thus, Disco predicts that BVCs should preserve their vector-coding properties in bowl-shaped geometries, but with shorter tuning distance compared with cylindrical environments (Paired t test: *t*_599_ = −20.93, P = 0.00, d = −0.85.) (Fig. 4, B to D).

**Fig. 4.**
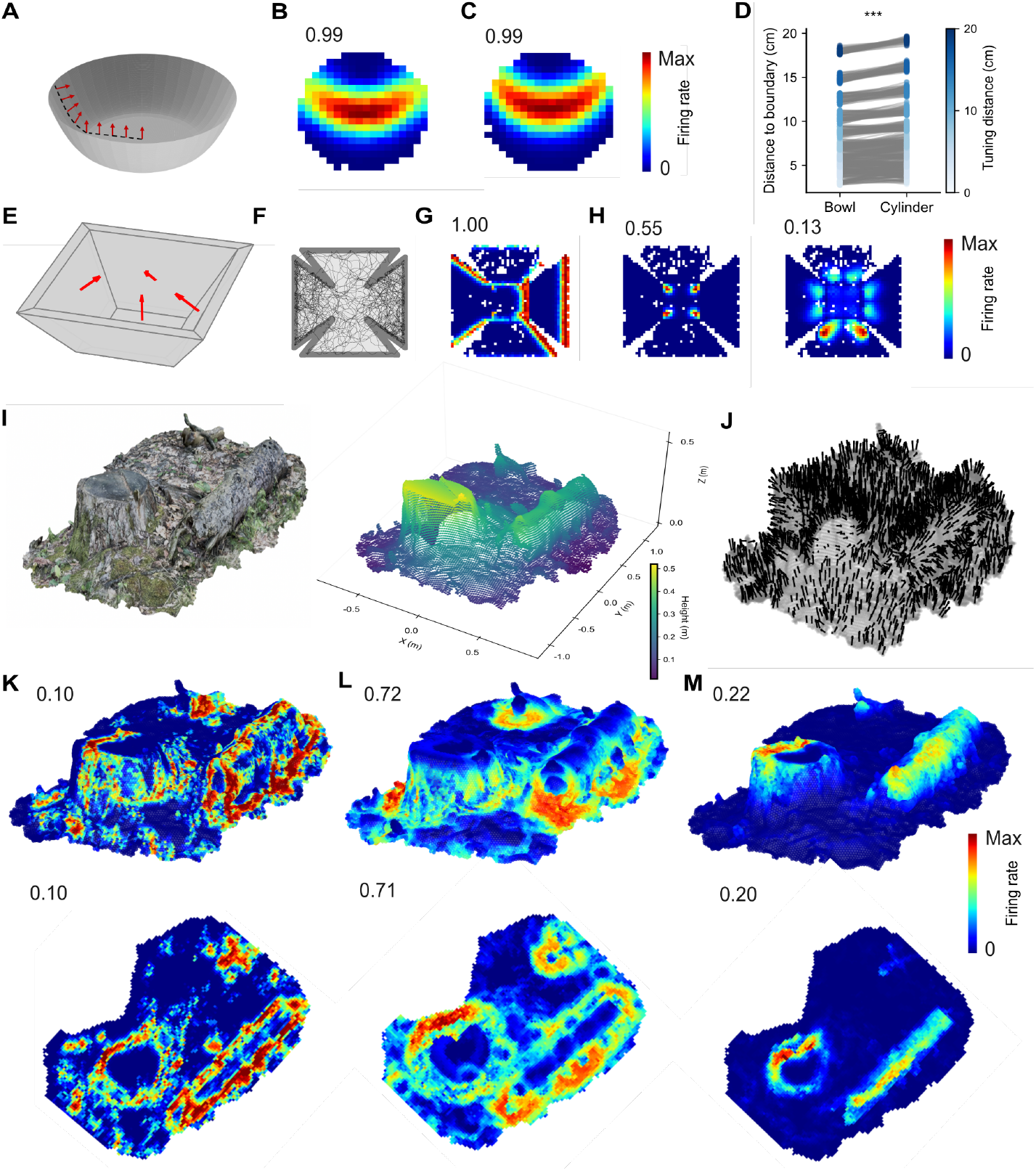
Predicted subicular activity in 3D laboratory and natural environments. (**A**) Continuous changes in surface normals (red arrows) in a bowl-like environment. (**B-C**) Rate maps of an example BVC in the bowl (B) and cylinder (C) environments. (**D**) Simulated BVC firing fields exhibit longer distances to environmental boundaries in the bowl compared with the cylinder. ***P < 0.001. (**E**) Surface normals (red arrows) of a sink-like environment with slanted surrounding walls. (**F**) Unfolded view of the agent’s trajectory. The agent can move across all surfaces of the sink environment, including the walls and the floor. (**G**) Rate map of an example BVC with the east tuning azimuth. (**H**) Two example corner cells with positive (left) and negative (right) tuning elevations. (**I**) A natural environment and the corresponding three-dimensional coordinates of surface pixels (*n* = 19129). (**J**) Surface normals (black arrows) for each surface pixel (gray points) computed from the coordinates (I) using a dimensionality-reduction method (materials and methods). Only 2000 surface normals are shown for visualization. (**K-M**) 3D rate map (top) and the corresponding bird’s-eye-view rate map (bottom). Rate maps were obtained by traversing all surface coordinates in (J) and are color-coded by firing rate (red, maximum; blue, minimum). The example BVC responds to a discontinuity located west of the agent (K). Two example corner cells respond to discontinuities at positive (L) and negative (M) tuning elevations relative to the agent. Image in (I) reprinted from BlenderKit (https://www.blenderkit.com) under a Royalty Free license.

Next, in the sink-environment, the agent could move along a horizontal floor and four surrounding inclined walls (Fig. 4E). Here, we modeled only surface-travelling navigation (Fig. 4F) and did not consider direction-unconstrained movement in volumetric spaces (e.g., air or water). Rate maps of Disco neurons form a mosaic of planar fragments (Fig. 4, G and H), consistent with previous work suggesting that the mammalian brain encodes 3D space in a quasi-planar manner (41). In addition to wall–floor intersections, Disco predicts that when an agent navigates along an inclined wall, BVCs also respond to wall–wall junctions, where discontinuities in surface normals are detected at fixed azimuths (Fig. 4G). That is, the transition from one inclined side of the sink to another has some of the character of a floor-wall transition, thus triggering BVCs. Moreover, corner cells respond to corners formed by two inclined walls during navigation on the floor, whereas cells with negative tuning elevations respond to corners formed by the floor and adjacent walls during navigation along walls (Fig. 4H).

To show that Disco can identify boundaries/corners in natural environments and predict corresponding subicular neural activity, we adopted a 3D natural scene from BlenderKit (https://www.blenderkit.com), which contains natural elements (e.g., tree trunks and grass) and provides full 3D coordinate information (Fig. 4I and fig. S12). Unlike the smooth planar surfaces typically used in laboratory settings, surface normals cannot be directly obtained. We applied eigenvalue decomposition to the local covariance matrix computed from the coordinates (materials and methods), and the surface normal at each point is defined as the eigenvector associated with the smallest eigenvalue (Fig. 4J). During locomotion on the ground, Disco BVCs and corner cells respond to upright boundaries (Fig. 4K) and corners (Fig. 4L) formed by the tree stump, stones etc. When the agent moves on logs or stumps, Disco BVCs respond to drops (Fig. 4K). Moreover, Disco predicts that corner cells with negative elevation tuning encode both the midline of cylindrical surfaces and basal edges. When the agent navigates near the central axis of a log or along the edge of a stump (Fig. 4M), these cells detect differences in surface normals on either side of the agent at a fixed elevation.

The high variance of geometric features in complex natural terrain raises the overall variability of neuronal activity. We suggest that a flexible internal gain setting (here implemented as thresholding) can regulate the activity of subicular cells to reduce noisy activations due to higher frequency changes in the terrain (supplementary text). We hypothesize that such a gain/threshold could be continuously adapted by the animal in a natural environment, in order to fine tune the coding resolution under noisy conditions (figs. S13 and S14).

### 2.5 Generalization to non-spatial domains

#### 2.5.1 BVC responses to discontinuities in color or texture space

Disco can generalize to non-geometric settings. Similar to experiments (21), we simulated an agent exploring an arena containing a traversable white stripe in the center (Fig. 5, A and B). The walls and floor were rendered in gray. In addition to wall responses derived from surface normals (Fig. 1A), model BVCs also fired at the stripe (Fig. 5C) by detecting a discontinuity in color space (or equivalently construed as texture space) (Fig. 5A). The underlying assumption is that BVCs can receive input from a range of modalities coded upstream. Disco computes a vector representation of the stripe by comparing lightness differences along a fixed Euclidean direction: the lightness values differ between the floor and the stripe (materials and methods). Previous studies have qualitatively classified the stripe as a sensory boundary without movement impediment (21). Disco quantitatively defines boundaries as discontinuities in either the spatial or non-spatial domain, allowing for BVC responses to walls, objects, drops, and stripes. This continues the theme that different subicular neurons are driven by varying and overlapping combinations of discontinuity information.

**Fig. 5.**
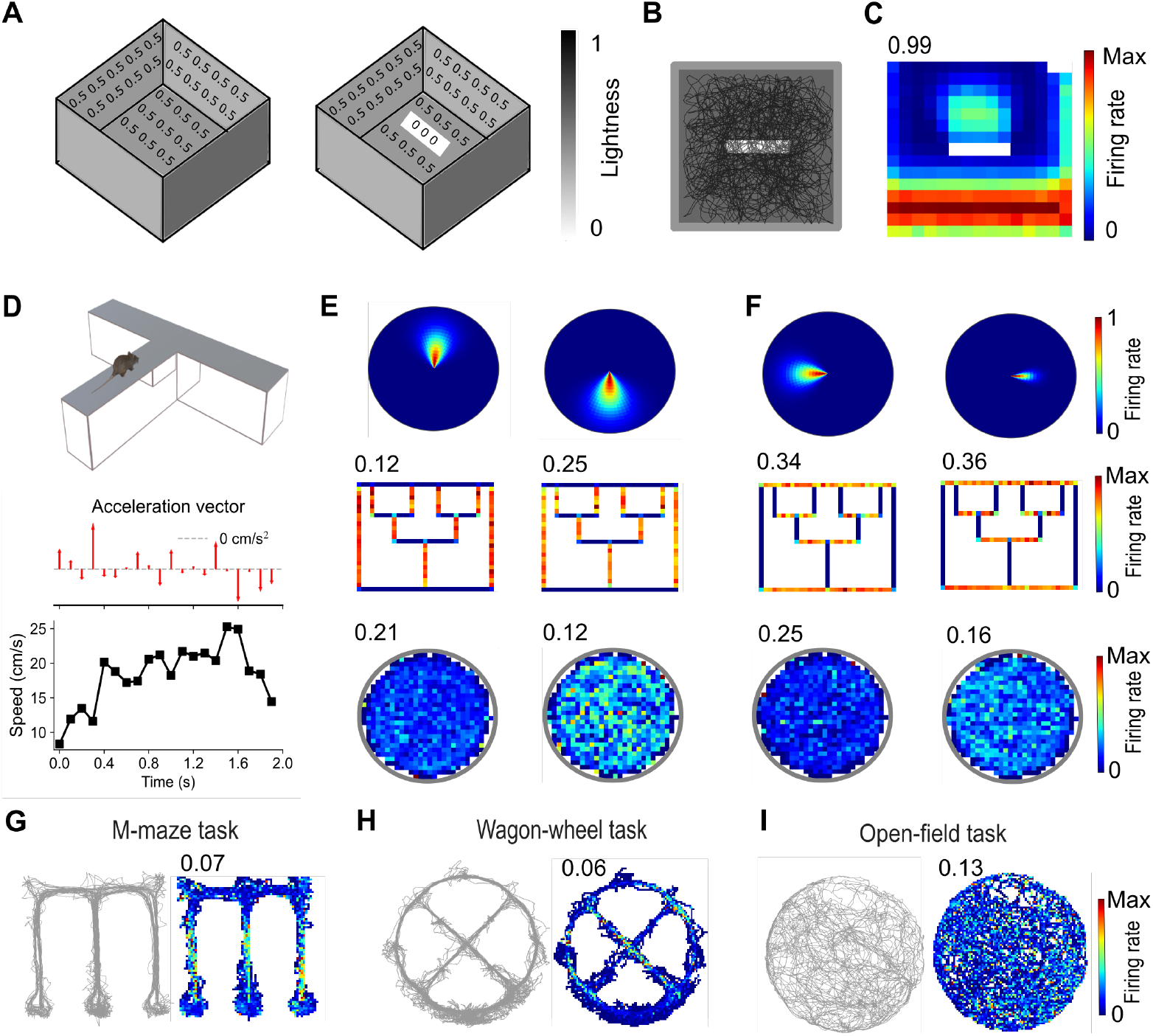
Extending the model to discontinuities in non-spatial domains. (**A**) A stripe is detected as a boundary through a discontinuity in surface color features, represented by differences in lightness values. (**B**) The agent’s trajectory can traverse the stripe. (**C**) Example rate map of a BVC responding to the stripe, corresponding to a discontinuity in color space. (**D**) Example acceleration and speed profiles of the agent during movement along a linear track. (**E**) Vector plots and rate maps of example axis-tuning cells selective for the y-axis. Top, the tuning vector is defined by the magnitude and direction of the acceleration vector. Example neurons respond maximally during acceleration (left) and deceleration (right) while moving along the positive y-axis. Middle, rate maps on the linear track. Bottom, rate maps in the open field. (**F**) Same as in (E), but for axis-tuning cells selective for the x-axis, responding maximally during acceleration (left) and deceleration (right) while moving along the negative x-axis. (**G-I**) Activity of simulated axis-tuning cells fitted to experimental trajectory data in linear-maze tasks(44) (G-H) and open-field environment(43) (I).

#### 2.5.2 Axis coding as discontinuities in behavioral state

Having explored combined geometric and non-geometric responses, we next asked whether Disco can be applied to purely non-spatial discontinuities, e.g., in behavioral state. Some subicular cells (axis-tuned neurons) encode the principal axes of an environment when rats run along a linear track (19) (Fig. 5D). BVCs could reproduce this axis coding by detecting the discontinuity in surface normals from the floor to the platform edge along the travel arm (fig. S15). However, many axis-tuned neurons exhibit irregular firing in open-field environments (19), suggesting their firing is not a mere variant of BVCs.

Inspired by the mixed selectivity BVCs that respond to stripes by detecting non-spatial (albeit localized) discontinuities, we posit that axis-tuned neurons respond to temporal discontinuities in movement speed, represented by the acceleration vectors (Fig. 5D). Using the same vector-coding base as for BVCs and corner cells, we modeled axis cells as tuning functions with fixed tuning acceleration magnitudes and angles (materials and methods).

We simulated an agent traversing a one-dimensional (1D) trajectory in the triple ‘T’ track maze, similar to experiments (19). For movement along the y-axis, Disco axis cells that maximally respond to deceleration or acceleration along the routes successfully encode the y-axis of the linear track (Fig. 5E). Cells tuned to acceleration in the positive direction respond to deceleration in the negative direction, and vice versa. Here we assume non-uniform velocity during unidirectional linear runs, producing both acceleration and deceleration events. This yields the prediction that fixing movement speed (e.g., by passively moving the animal at a constant speed in a cart) abolishes axis coding. Corresponding results were also observed for movement along the x-axis (Fig. 5F). The irregular firing fields in the open field can be attributed to the random exploratory trajectories (Fig. 5, E and F). To further verify our proposed mechanism, we fit Disco axis cells to the published datasets of animals’ trajectories from both and open-field (43) and linear-track (44) tasks (Fig. 5, G to I, figs. S16 and S17). Disco axis cells exhibit axis tuning in the M-maze and wagon-wheel maze, and reproduce the irregular firing patterns observed in the circular arena.

### 2.6 Event boundaries as discontinuities in neural population activity

The above results show that Disco can account for a wide range of single-cell responses in the subiculum within a single framework. Next, we asked whether the core principle can be applied to the population level. To visualize how environments are represented in a low-dimensional neural manifold, we calculated 2D embeddings (45) of the population activity of all simulated Disco neurons with geometric codes (boundary and corner vectors) in two-compartment environments (materials and methods). The two compartments were connected either by a corridor, by a doorway, or directly attached (Fig. 6, A and C). Consistent with a previous study (3), Disco showed that the neural manifold in the subiculum can distinguish environments with different geometric shapes. We found that two identical compartments showed largely overlapping representations (Fig. 6B). Differing shapes produced clearly separable representations (Fig. 6D), and this distinction was insensitive to object insertion. We hypothesize that incorporating non-geometric discontinuities (Fig. 5), guided by experimental data, could further disambiguate population codes between similarly shaped environments.

**Fig. 6.**
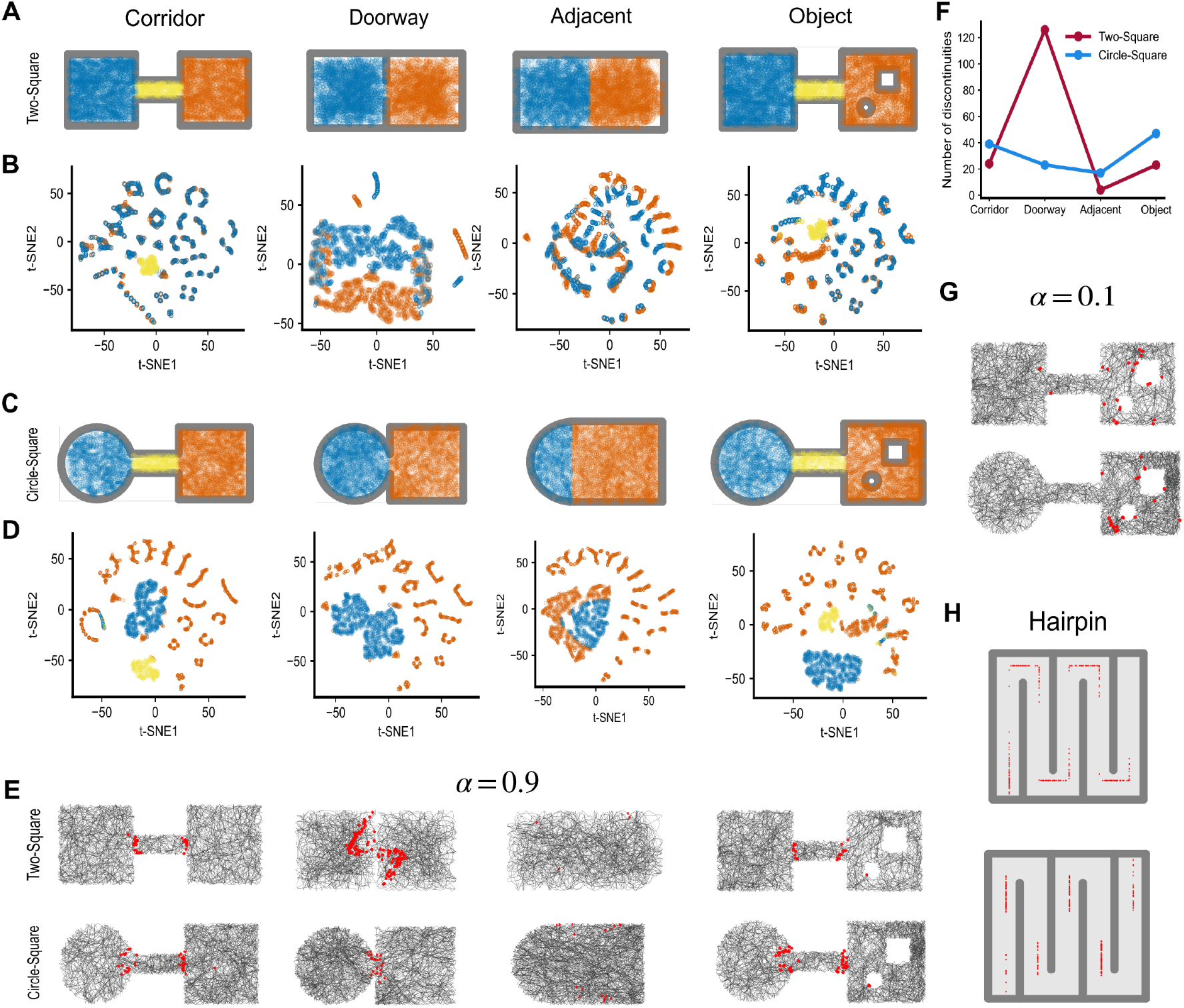
Temporal discontinuities in population activity. (**A**) Simulated positions in arenas formed by two squares connected in different configurations. (**B**) Two-dimensional embedding (t-SNE) of population activity in the two-square arenas for non-corner cells, corner cells, and BVCs. Each dot represents the population state at a single position. Arena regions of agent positions are color-coded as in (A). (**C-D**) Same as (A-B), for arenas formed by a square and a circle. (**E**) Temporal discontinuities in population activity signaling event boundaries during exploration. The agent’s trajectory is shown in black. Red dots indicate time points at which the difference in cumulative population activity (non-corner cells, corner cells, and BVCs), with a large discount factor (α = 0.9), exceeds 0.1. (**F**) Number of temporal discontinuities identified in each environment. (**G**) Same as (E), for temporal discontinuities corresponding to cumulative population activity computed with a small discount factor (α = 0.1). (**H**) Temporal discontinuities in the linear-track task, identified from population activity including axis-tuned cells (top) or excluding axis-tuned cells (bottom).

We next examined whether Disco can capture contextual changes. Dimensionality reduction techniques, while illustrative, do not speak to dynamical mechanisms. To obtain an actionable measure of contextual changes, we constructed a population vector at each time point, composed of the discounted average accumulated firing rate of all Disco neurons. This yields a memory signal of exploratory experience. Discontinuities in this neural space were defined as time points at which the cosine distance between adjacent population vectors exceeded a threshold (materials and methods). In the corridor-connected environments, the discontinuities occurred predominantly when the agent transitioned between a compartment and the corridor (Fig. 6E). We also found that discontinuities were detected near the doorway, and were greatly reduced once the doorway was removed (Fig. 6, E and F), consistent with previous experimental findings (46). Experiences of the world are thought to be segmented into successive discrete events that can later be reconstructed from memory (24,47–49). Here, we interpret discontinuities in the population code as event boundaries (i.e., transitions in the exploration context), a signal that could for instance trigger remapping in the hippocampus (50–54).

We also found that object insertion did not significantly alter Disco’s ability to detect contextual changes (Fig. 6, E and F). However, using a smaller discount factor led to event boundaries appearing near objects (Fig. 6G). This suggests the subiculum could encode event boundaries using in a distributed mechanism, potentially subject to neuromodulatory signals (55), with rapid memory updating for local features (e.g., object novelty) and slower integration for global contextual changes. Alternatively, local computation within subiculum could read out this population change, with specific subicular cells signaling temporal discontinuities.

Disco axis cells did not significantly affect our manifold analysis or event-boundary identification (figs. S18-S19). This likely reflects the irregularity of axis-cell firing in open fields (19) (Fig. 5, E, F and I), as well as limitations in our simulated trajectories, which lack some behavioral details (e.g., thigmotaxis (56)). However, behavioral changes are one contributor to event boundaries (48,54). We simulated a virtual rat running a linear trajectory in a hairpin maze (Fig. 6H), where Disco axis cells exhibit systematic firing patterns. We found that incorporating axis-tuned signals caused discontinuities to cluster at the turning points, providing a more comprehensive representation of event boundaries. This signal could also trigger re-anchoring of grid cells to turns in the hairpin maze, for which individual track segments have previously been fitted discontinuous 1D slices through 2D grids (57).

## 3 Discussion

Accounting for seemingly disparate phenomena under the umbrella of a common organizing principle is one of the key goals of scientific theories. Disco provides such a framework in which BVCs and corner cells are represented in a vector format with complementary 3D receptive fields, offering a common explanation for their selectivity to diverse local geometric features (e.g., walls, drops, objects and corners), driven by discontinuities in the spatial domain (Figs. 1 to 3). Disco Corner cells reproduce fine-grained experimental observations (including height-dependence) in convex, concave and continuous flavors, and suggest some nominally non-corner cells are driven by the same discontinuity information as corner cells. We predict subicular activities in 3D space and show robust applicability to natural environments (Fig. 4). In non-spatial domains: discontinuities in color/texture space can account for BVC responses to stripes (framed as mixed-modality coding), and discontinuities in behavioral space explain axis-tuned neurons (Fig. 5). At the population level, Disco enables discrimination between environments of different shapes, and the extracted temporal discontinuities can signal event boundaries (Fig. 6).

The question remains, is discontinuity detection just a convenient algorithmic proxy or a signature of a ground-truth neural coding principle? One prior study has used optic flow templates computed from surface normals to drive BVC activity (26). However, this study did not explore the use of discontinuities as a generalized coding principle. Here we propose that the coding of discontinuities in different combinations may be a key feature of subicular activity, strongly supported by the fact that Disco reproduces numerous single-cell phenomena with fine granularity (e.g., the reproduction of discrete and continuous corner cells (3)); the applicability to natural environments; and purely non-spatial settings like behavioral state and event boundaries.

### 3.1 Relationship to anatomy

The diversity of single-cell tunings suggests that information about environmental features is computed upstream of the subiculum through the integration of multiple information sources. This was exemplified in our simulation of BVCs responding to a stripe on the floor, where it was suggested that BVCs integrate information from multiple modalities. Targets for investigation of this hypothesis should be the known subicular input pathways (9,11,58,59). Axis cells, coding for acceleration, may rely on path-integration signals from medial entorhinal cortex (MEC), where speed cells (60,61) have also been identified. Alternatively, optic flow signal could contribute. Several of the connections between the subiculum and these regions (including MEC) are reciprocal (11,62,63). Grid-cell remapping at movement turning points (64) may be supported by event-boundary signals facilitated by the population code with axis cells (Fig. 6h). Potentially Disco-like mechanisms could also produce spatial responses reported in other areas (6,65), as is suggested in the supplementary text for (egocentric) vertex cells (RSC (35)).

Previous work has shown connectivity between the subiculum and visual cortex (66–68), which carries neural representations of color and orientation (22,69) and may contribute to identifying discontinuities. Similarly, connections with the anterior thalamic nucleus (58) potentially place the subiculum only one synapse away from RSC, limbic and sensory areas (70). Spatial coding is mainly associated with the dorsal subiculum, while the ventral subiculum – connected with the limbic system and prefrontal cortex – is implicated in motivational and affective regulation. The subiculum has been shown to encode variables of decision-making and navigation tasks (71,72) (e.g., reward and trajectory). Disco could be integrated with an inter- or intra-subicular network model (34,38) to further study cognitive functions and behaviors (e.g., proximodistal CA1–subiculum pathways for object and spatial processing (9,38,62,73,74)). Some cell types identified in the subiculum (e.g., place cells (19,75)) could result from local interactions. Thus, to complement Disco, we also added a circuit-level model, suggesting how intra-subicular network interactions could contribute to the neural responses (figs. S8 to S11). Moreover, both spatial and non-spatial discontinuity coding may influence the recipients of subicular projections (9,76,77), such as event-boundary representations in the lateral entorhinal cortex (48), arousal regulation via the locus coeruleus (78), and novelty detection in the striatum (79). Sudden onset/change of affective signals could also be construed as a discontinuity.

### 3.2 Predictions and related work

Several experimentally testable predictions can be derived. For instance, BVCs should respond to holes; corner cells fire at corners identified from the animal’s top-down perspective; and axis-tuned neurons could exhibit structured firing patterns during wall-following behavior in open fields. Extrapolating, Disco could also be expanded to predict neural activity across diverse environmental configurations and manipulations, for example: the evolution of subicular activity in morphed mazes (80) or rats navigating through burrow systems. The present model helps clarify several previously ambiguous findings, including the definition of boundaries (5,81); why some BVCs respond predominantly to platform edges whereas others are tuned to the edge of reachable space (5); the (non-)overlap between axis-tuned neurons and BVCs (19); the classification of corner cells, and how different types of event boundaries (24,48) (e.g., contextual changes versus novelty detection) can be distinguished via population dynamics.

Finally, the theory may motivate a subtly reframed view of hippocampal memory with regard to indexing and associative processing (27,28). Given the conserved hippocampal anatomy across species, we may hypothesize that the hippocampal system performs similar computations while operating on different sensory inputs specific to species. Disco is highly compatible with this notion by proposing that the hippocampus binds discontinuities of experience (e.g., encountering an object, the onset of a smell, a sudden startle) but not the items per se. Upon retrieval, representations of these discontinuities must be unpacked to retrieve item representation via cortical pattern completion (27,30,85). The privileged anatomical position of the subiculum supports this process by routing discontinuity signals to distinct brain areas (71) and onwards via a cortical cascade of reinstatement (30). How far back the cascade reaches depends on the information required by the remembering agent (e.g., the level of conceptual or sensory detail dictated by task demands). The reconstructed representations could be sensory, motoric, conceptual, or affective, according to what modalities were engaged during memory encoding.

## 4 Conclusion

The subiculum is the main output layer of hippocampal formation, the latter being long regarded as the neural substrate of “cognitive map” theory (86,87). Grid cells in the medial entorhinal cortex provide a context-independent spatial metric and directional information, bridging hippocampal place-field representations with physical spatial coordinates via path integration. The subiculum can exhibit context-independent encoding of environmental features. Individual neurons display tuning to a multitude of elements that are highly abstract and do not readily admit explanations based on any single sensory modality. Disco offers a common explanation for both geometric and non-geometric features and supports a broader conceptualization of cognitive maps.

Potentially, discontinuity detection can be extrapolated as a computational principle to other brain areas. It bares resemblance to novelty detection (79) but is not identical: an environmental element can be a discontinuity without being novel. However, sudden changes in stimulus attributes (novel or not) can be signaled by one of the most fundamental neural properties: spike-frequency adaptation (88). The uncertainty about the precise location of the discontinuity, as expressed through the gaussian vector base, endows cells with properties resembling classical feature selectivity (e.g., orientation selectivity in the early visual system (22)). But what is detected is the change in sensory attributes (or internal states). Disco suggests that such a simple and transparent framework could operate far from the sensory periphery and serve as general coding principle in the subiculum, consistent with its role in spatial and non-spatial coding.

## 5 Materials and Methods

Reference (17) introduced a BVC tuning function that uses two Gaussian components to model neural responses as functions of vector distance and angle. This formulation can reproduce the firing-rate patterns of individual neurons across diverse environments. However, it requires a priori knowledge of the boundary’s location and the specification of boundary/object status of an environmental feature. Empirically it is observed that BVCs can simultaneously respond to barriers, objects, edges, and stripes (5,21,31).

In our algorithm, beyond the Gaussian components used for vector coding, the tuning function of BVCs includes a discontinuity detector that we interpret as the output of upstream brain regions. This detector identifies boundaries/drops/objects via 3D receptive fields (Fig. 1), by detecting salient changes in surface normals along the vertical axis. By detecting salient changes horizontally, we also formulate corner-cell activity within a similar vector-coding framework, as detailed below. That is, the same coding principle applies with a different mix of detectors. This is principal element of variation among cell types. Most simulations were based on 2D movement trajectories generated using RatInABox (56). Simulation details are provided in the supplementary text. We also extend the model to the natural environment where the agent navigates in 3D space.

We then show that our model can generalize to non-geometric space. In particular, by detecting discontinuities in velocity we obtain axis cells, which implement vector coding for acceleration. Finally, by analyzing population activity in the subiculum, we demonstrate that temporal discontinuities in neural space can signal the corresponding event boundaries. All model parameters used in the simulations presented in the main text are listed in table S1. Below we further outline possible circuit level interactions and compare pre-existing models, specifically with regard to corner-coding.

### 5.1 Discontinuities in Euclidean space

We simulated an agent exploring environments with various geometric features (fig. S1 and table S2). At time *t*, each 3D point in the environment is expressed in spherical coordinates (*l, θ, φ*) relative to the agent’s body centroid (*x*_*o*_(*t*) ∈ ℝ^3^). The key elements contributing to the firing of BVCs and corner cells are the normal vectors of perceptual surfaces. Then each point is associated with a feature value *f*_*t*_(*l, θ, φ*), representing the normal vector of the surface (e.g., wall, floor) at that location. For points located outside any surface (e.g., in air or underground), *f*_*t*_(*l, θ, φ*) was undefined.

Along each direction (*θ, φ*), we define a detection vector *V*_*t*_(*θ, φ*) originating from *x*_*o*_(*t*) to the first intersection with an environmental surface. The length of this vector, *l*_*t*_(*θ, φ*), represents the radial distance at which the feature value changes. If no such discontinuity in *f*_*t*_(*l, θ, φ*) occurs within the interval *l* ∈ (0, *L*], the vector distance is set to *l*_*t*_(*θ, φ*) = *L*. The numerical step size along the radial dimension was *dl*.

Here, we assume the detection vector’s feature values *f*_*t*_(*l*_*t*_(*θ, φ*), *θ, φ*) determines the activities in subiculum. A discontinuity at direction (*θ, φ*) can drive the firing of different cell types. The vertical and horizontal discontinuity indicators, restricted to jump discontinuities, are given by

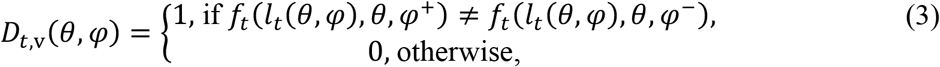

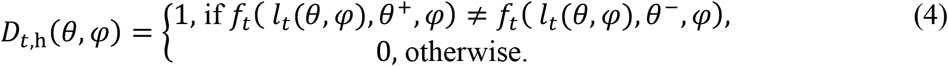

The numerical elevation and azimuth used for simulation are *dφ* and *dθ*.

#### Definition of a BVC

Unlike the previous 2D receptive fields, we redefined BVCs’ receptive fields in a 3D spherical coordinate system, allowing for a more general setting of tuning functions (see below). The receptive field is responsible for detecting discontinuities based on directly received sensory input (*f*_*t*_(*l*_*t*_(*θ, φ*), *θ, φ*) . BVCs are defined as neurons responding to the vertical discontinuity indicated by *D*_*t,v*_(*θ, φ*). For example, a segmentation between the wall and the floor is present along this direction at a distance *l*_*t*_(*θ, φ*) . For each azimuth *θ*, the discontinuity associated with the shortest detection vector (e.g., pointing to the nearest wall) contributes to neural activity.

#### The tuning function of BVCs

The BVC tuning function is composed of two Gaussian components responding to the vector distance (*l*_*t*_(*θ, φ*)) and horizontal angle (*θ*), and the discontinuity indicator *b*_*t*_(*θ, φ*) = *D*_*t,v*_(*θ, φ*). The vertical discontinuity contributes to the firing rate of a BVC unit *k* (tuned to distance *d*_*k*_, azimuth *ϑ*_*k*_) as follows,

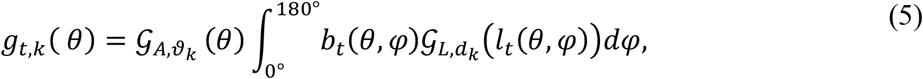

where an elevation of 180^◦^indicates directly upward and 0^◦^indicates downward. 𝒢_*L*_ is a Gaussian radial function, which is given by

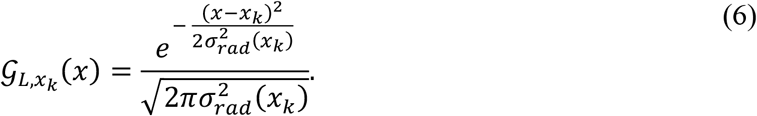

𝒢_*A*_ is the radial von Mises distribution (a generalization of a Gaussian for periodic variables):

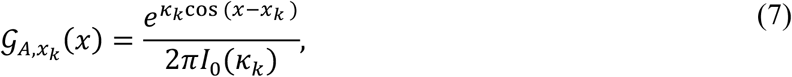

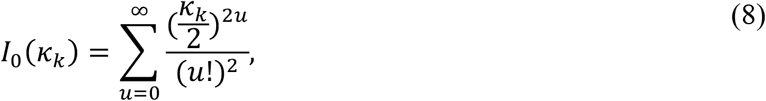

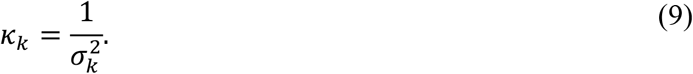

In our simulation (Figs. 2, 4, 5B-C, 6), the tuning azimuths were uniformly distributed with a spacing of 6^◦^, *ϑ*_*k*_ ∼ 𝒰[0◦, 360◦]. The tuning distances *d*_*k*_ increased linearly from 2.97 cm to 19 cm over a radial extent of 15 distance units. The radial tuning width *σ*_*rad*_(*d*_*k*_) scales linearly with the tuning distance, 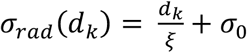, with *ξ* and *σ*_0_ denoting fixed constants. Across all simulated neurons in our study (see below; BVCs, axis cells, and (non-) corner cells), the angular width *σ*_*k*_ is drawn from a uniform distribution, *σ*_*k*_ ∼ 𝒰[10◦, 30◦].

At time *t*, the contribution of all detected discontinuities to the firing of the unit *k* is determined by integrating Equation. (5) over *θ*:

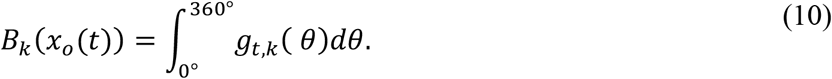

Here, a bearing of 0^◦^corresponds to east, 180^◦^ to west, and bearings of 90^◦^and 270^◦^to north and south, respectively.

#### The two-point model for BVC responding to drops

To explain the experimental observation that BVCs’ rate maps extend beyond the platform edges (5), we extend our model to a two-point formulation. The agent is represented by two points, the head and the body centroid, connected by a 5 cm rigid link. In this variant, we assume a functional dissociation between discontinuity detection and neuronal tuning. Discontinuities are detected based on feature values sampled at the head position (*x*_*h*_(*t*) ∈ ℝ^3^), but the tuning vector of each BVC is defined with respect to the body centroid (*x*_*o*_(*t*)), consistent with the original formulation.

Accordingly, the detection vector, *V*_*t*_(*θ*^*h*^, *φ*), is redefined to originate from *x*_*h*_(*t*). *θ*^*h*^ is the azimuth of the discontinuity relative to the head. To compute the tuning vectors that facilitate the firing of BVCs, a coordinate transformation is applied following the detection of a vertical discontinuity (Equation. (3)). The tuning vector is given by

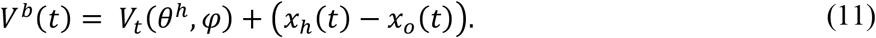

Then, we compute the distance 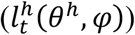 and azimuth (*θ*^*b*^) of the discontinuity relative to the body centroid.

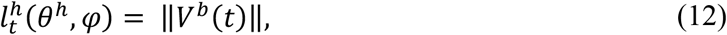

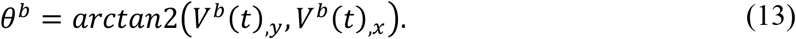

The contribution of the detected discontinuity to the neuron *k* is defined as follows,

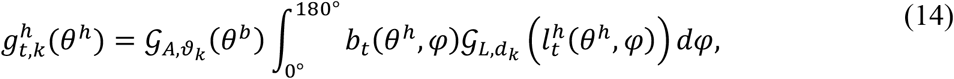

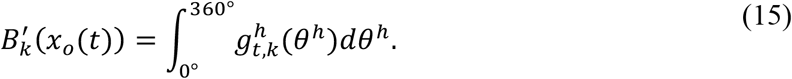

and the firing rate is determined by detecting discontinuities across the angular range of head point*θ*^*h*^:

This two-point model is only used for the analysis of BVCs’ responses to the drops (Fig.2 F to I). All other simulations employ the original centroid-centered formulation.

#### The egocentric BVC

We also reproduced the features of egocentric BVC activities by reformulating Equation. (5). The detector, *D*_*t,v*_(*θ, φ*), determines the allocentric bearing of discontinuities. The egocentric bearing was then calculated by subtracting the agent’s allocentric head direction, which is given by:

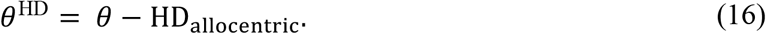

The contribution of the vertical discontinuity to the firing rate of neuron *k* is given by:

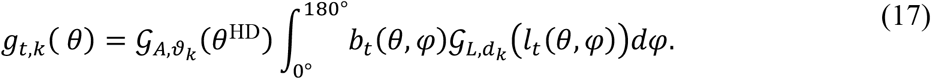

An egocentric bearing of 0◦ corresponds to a discontinuity in front of the agent, and 180◦ denotes one located behind the agent. Bearings of 90◦ and 270◦ represent directions to the left and right of the agent, respectively.

#### Definition of corner cell

To the best of our knowledge, this is the first proposal of a dedicated tuning function of the corner cell (3), which has format consistent with BVCs. One prior model proposes vertex coding (without convexity/concavity classification) as emerging from the superposition of BVCs (35). The corner is defined by three factors: wall height, wall connectivity and the angle between two walls. Our sampling mechanism explains how “wall height” and “wall connectivity” affect the firing rate of the corner cell. Additionally, “convexity/concavity” coding can be accounted for by manipulating normal vectors to control the firing of the corner cells.

We modelled corner cells using the same receptive field structure as for BVCs. We first simulated neurons responding to horizontal discontinuities, which can be detected by *D*_*t,h*_(*θ, φ*). Then, neurons with positive corner scores (cornerscore_cell_ > 0, see below) were classified as corner cells, whereas all remaining cells were classified as non-corner cells. For example, a segmentation between two adjacent walls is present along this direction at a distance *l*_*t*_(*θ, φ*). At the fixed elevation *φ*, the discontinuity associated with the shortest detection vector (e.g., pointing to the nearest corner) contributes to neural activity.

#### The tuning function of corner cells

The function includes two Gaussian components responding to the vector distance (*l*_*t*_(*θ, φ*)) and elevation angle (*φ*), and the discontinuity indicators *c*_*t*_(*θ, φ*) = *D*_*t,h*_(*θ, φ*). The horizontal discontinuity contributes to the firing rate of neurons *k* (tuned to distance *r*_*k*_, elevation *φ*_*k*_), is given by

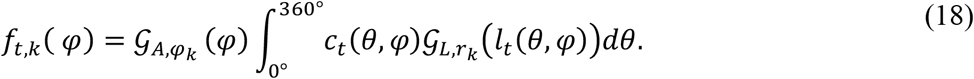

In our simulation (Fig. 3, A to H), the tuning distances *r*_*k*_ increased linearly from 0.73 cm to 24.5 cm over a radial extent of 15 distance units. The tuning elevations were uniformly distributed with a spacing of 6^◦^, *φ*_*k*_ ∼ 𝒰[0◦, 180◦].

We used a semicircular von Mises tuning function model the elevation tuning of neurons. At time *t*, the contribution of all corners to the firing of the neuron *k* is determined by integrating Equation. (18) over *φ*:

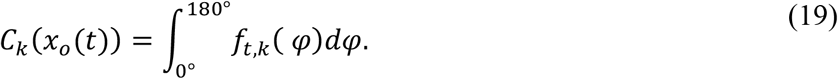

#### Effects of angles on corner cells

Experimental evidence showed that more acute corner angles elicit higher firing rates (3). We model this by calculating the dot product between the two neighboring normal vectors. We define

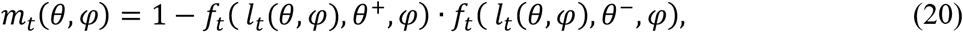

which is bounded within the interval [0,2]. Larger values of *m*_*t*_(*θ, φ*) correspond to larger angles between the surface normals, which in turn indicate sharper corners. When *m*_*t*_(*θ, ϕ*) = 0, two normal vectors are identical, indicating the absence of a corner. Then, the binary discontinuity indicator *D*_*t,h*_(*θ, φ*) can be formulated as a continuous-valued measure as follows,

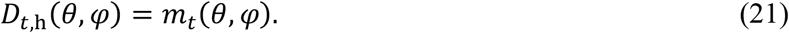

#### Effects of convexity on corner cells

To model the observation that corner cells can be divided into convex and concave populations (3), we extended the elevation range from a semicircular (*φ*) to a full circular domain. The angular space was then partitioned into two non-overlapping semicircular ranges: [0◦, 180◦] for concave tuning and [180◦, 360◦] for convex tuning. Algorithmically, the convexity is identified by computing the cross product between two adjacent normal vectors.

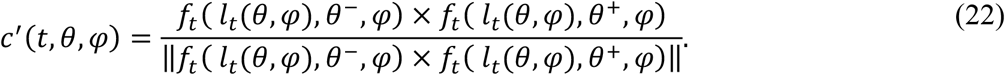

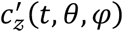 is the directional indicator, which captures the orientation of the cross product along the final coordinate axis. Here, 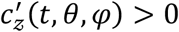 corresponds to the concave corner, while 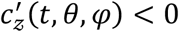 indicates to the convex corner. We define the discontinuity indicators with effect of convexity as

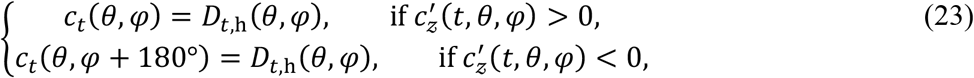

which distinguishes concave and convex corner configurations by assigning antipodal tuning angles. The firing rates of neurons with concave (*k*_*concave*_) and convex (*k*_*convex*_) tuning are defined as

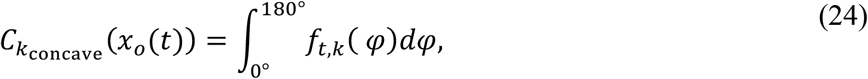

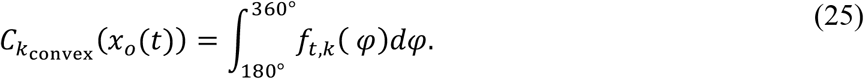

After including the effects of convexity, in our simulation (Figs. 3I to P, 4, 6), the tuning elevations were uniformly distributed with a spacing of 6^◦^, *φ*_*k*_ ∼ 𝒰[0◦, 360◦] . The tuning distances increased linearly from 1.12 cm to 14.5 cm over a radial extent of 15 distance units.

### 5.2 Non-spatial discontinuities

#### BVC responses to discontinuities in color/texture space

BVCs can respond to the white stripe in the environment with gray surfaces (21). We assumed the white stripe as a discontinuity in color space (alternatively texture space).

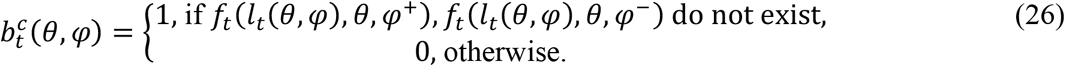

Here, *f*_*t*_(*r, θ, ϕ*) ∈ ℝ represents the lightness value of the surface at that location. Here, the white stripe has a lightness of 1, and the gray walls and floor had a lightness of 0.5. For points located outside any surface (e.g., in air or underground), *f*_*t*_(*l, θ, φ*) was undefined.

The contribution of the color discontinuity to the firing rate of neuron *k* is given by:

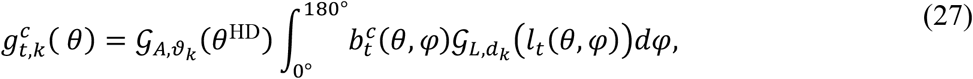

and the firing rate is determined by discontinuities in both Euclidean (Equation. (5)) and color space, which is defined as follows,

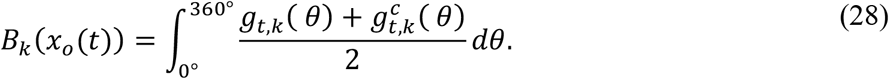

#### Axis cell responses to discontinuities in behavioral space

Within the Disco framework, we predict that the recorded axis-tuned neurons (19) are tuned to discontinuities in movement speed. The corresponding discontinuity indicator is defined as

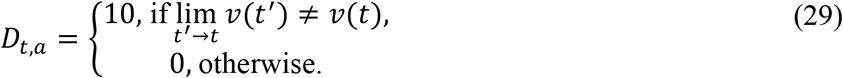

*v*(*t*) denotes the agent’s velocity at time *t*, which is given by

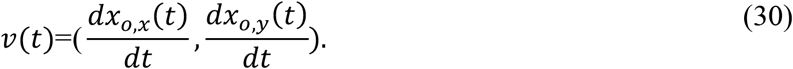

Here, we consider only 2D trajectories and use the *x*- and *y*-components of *x*_*o*_.

#### Definition of axis cells

Axis cells are defined as neurons responding to the discontinuity indicated by *D*_*t,a*_. The firing of each neuron is assumed to represent the acceleration vector, which is given by

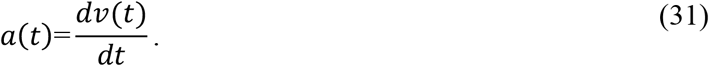

#### The tuning function of axis cells

Consistent with other subicular neurons, we model axis cells in a vector-based format. The function is composed of two Gaussian components that respond to the magnitude (*l*_*a*_(*t*)) and direction (*θ*_*a*_(*t*)) of acceleration.

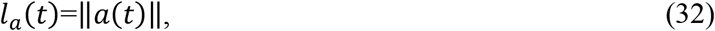

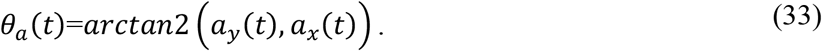

At time point t, the firing rate of unit *k* (tuned to magnitude *d*_*a,k*_, direction *ϑ*_*a,k*_), is defined as

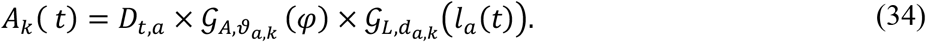

In our simulation (Figs. 5E to I, 6), the tuning directions were uniformly distributed with a spacing of 6^◦^, *ϑ*_*a,k*_ ∼ 𝒰[0◦, 360◦]. The tuning magnitudes *d*_*a,k*_ increased linearly from 1.48 cm/s2 to 9.5 cm/s2 over a radial extent of 10 distance units.

### 5.3 Discontinuities at the population level

For each time point *t*, neural population activity was represented by a population vector, M(*t*), which is given by

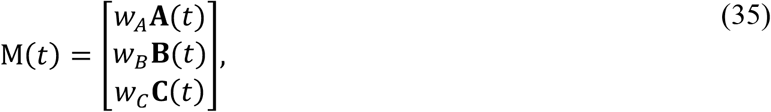

where 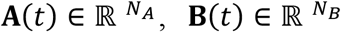 and 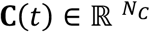 denote the firing-rate vectors of the three types of simulated neurons. Here, *N*_*A*_, *N*_*B*_, and *N*_*C*_ are the numbers of the simulated Disco axis cells, BVCs, and corner and non-corner cells. *w*_*A*_, *w*_*B*_, and *w*_*C*_ are scalars scaling the relative contributions of each neuron type to the population activity. When Disco axis cells’ activities were taken into account in the analysis (Fig. 6H), *w*_*A*_ = 100; otherwise, default parameter values were used (table S1).

#### Event boundaries represented by temporal discontinuities

We defined the encoding of experience in population activity as a discounted accumulation of activity, computed recursively as

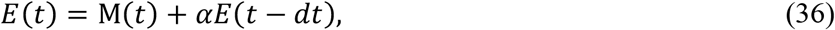

where *α* ∈ (0,1) is a discount factor controlling the influence of past memory (Fig. 6). To identify discontinuities in experience, we computed the cosine distance between adjacent time points.

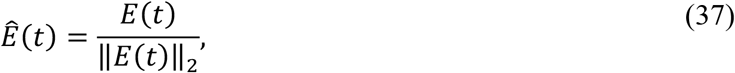

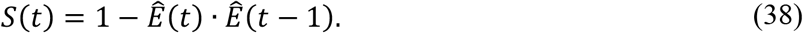

The discontinuity indicator is defined as

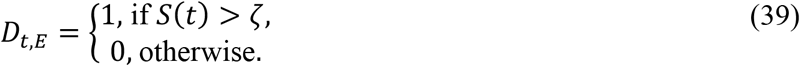

A discontinuity is detected at time *t* when the distance between successive vectors falls above the threshold *ζ*, which we interpret as an event boundary. *ζ* was set to the 90th percentile of the distance distribution for the linear-track simulation (Fig. 6H), 99.8th percentile for the open-field simulation with *α* = 0.1 (Fig. 6G). To enable comparison of discontinuity counts across different open-field simulations with *α* = 0.9 (Fig. 6, E and F), *ζ* was set to the 99.8th percentile identified in the two-square compartments connected by a corridor as a reference.

### 5.4 Analysis

#### Rate map

For simulations in laboratory settings, rate maps were obtained by discretizing the environment into 2 × 2cm spatial bins and computing the average firing rate for each bin based on activity recorded when the agent occupied that location.

For the simulation in natural environment, 3D rate maps were generated directly by computing firing rates at surface points represented by their 3D coordinates. Bird’s-eye-view rate maps were generated by projecting the 3D surface coordinates onto the 2D (X, Y) plane, followed by the same computation procedure used for laboratory settings.

#### Egocentric boundary rate map

Data were analyzed in an egocentric reference frame, where the boundary location was defined relative to a static agent position. Here, the boundary locations were represented by the vector (*l*_*t*_(*θ, φ*), *θ*) obtained from our proposed discontinuity indicator *D*_*t,v*_(*θ, φ*). The main components used are the agent’s head direction and the position of detected discontinuities relative to the agent. The 360° surrounding the agent was divided into angular bins of 10°, centered at 0° along the current head direction. Radial distance from the agent’s position was discretized into 1 cm distance bins equal to half the length of the longest wall, yielding a polar grid with 10° × 1 cm bins. Egocentric boundary rate maps were obtained by computing the average firing rate for each bin based on activity recorded when a discontinuity was present within that bin.

#### Spatial information

To quantify the information content of a neuron’s activity, we calculated spatial information in bits per spike (89) for each neuron as follows,

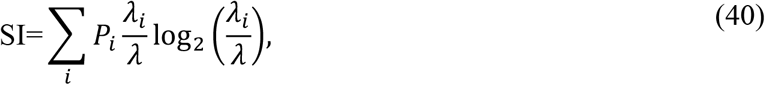

where *P*_*i*_ is the probability of the agent occupying the *i*th spatial bin (5×5cm), *λ*_*i*_ is the neuron’s average firing rate in the *i*th bin, and *λ* is the average rate across the entire simulation.

#### Peak firing rate

The peak firing rate was defined as the maximum firing rate across all bins in the rate map.

#### Corner score

We defined corner score following the reference (3). Place fields were first identified as contiguous regions of the 2D rate map with firing rates exceeded 0.3 times the peak firing rate. For each field, the corner score was defined as:

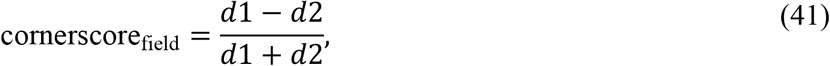

where *d*1 is the distance between the bin with the maximum firing rate within the field and the centroid of the arena, and *d*2 is the distance between that bin and the nearest corner. The score ranges from −1 for fields located at the centroid to 1 for fields located precisely at a corner.

In an environment with *k* corners, the corner score of a cell with *n* fields is defined as:

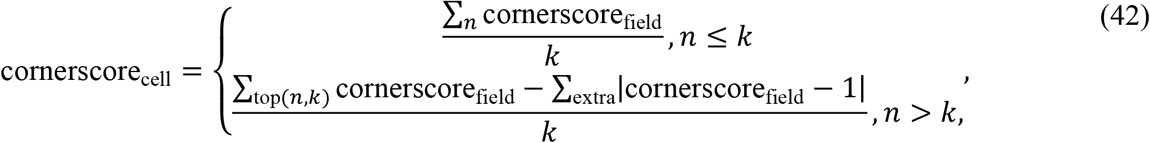

where top(*n, k*) refers to the *k* fields with the highest corner scores among the *n* fields. extra denotes the corner scores of the remaining fields and acts as a penalty term for cells with excessive fields.

#### Surface normal estimation in 3D point clouds

For the natural environment, we used a free asset sourced from BlenderKit (https://www.blenderkit.com), allowing direct access to a point cloud, **x**_*i*_ ∈ ℝ^3^, *i* = 1, …, 19129, which represents points sampled from the surface of the environment. Each point is defined by its 3D coordinates.

For each point **x**_*i*_, the normal was computed analytically based on its *k*_*n*_-nearest neighbors in Euclidean space, 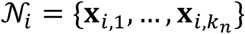. Here, we set *k*_*n*_ = 30. The neighborhood points were first centered by subtracting their centroid.

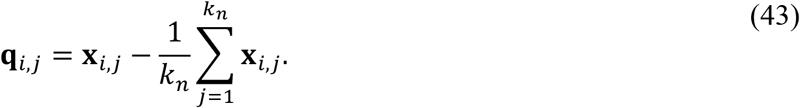

The surface normal (**n**_*i*_) at point **x**_*i*_ was determined from the local covariance matrix (**H**_*i*_) via eigenvalue decomposition.

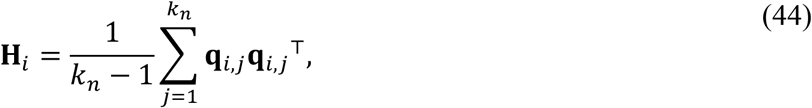

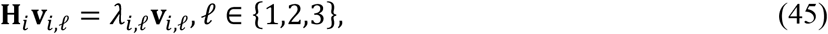

where the eigenvalues were ordered as *λ*_*i*,1_ ≤ *λ*_*i*,2_ ≤ *λ*_*i*,3_ . The eigenvector **v**_*i*,1_ represents the direction of minimal variance in the local point cloud. To obtain the surface normal, **v**_*i*,1_ was normalized and orientation-corrected with respect to a reference direction u = (0,0,1)^⊤^.

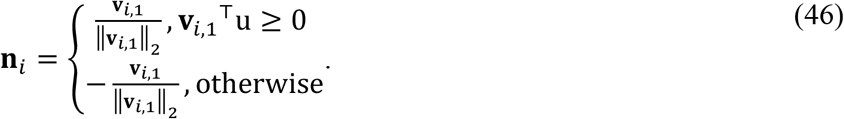

#### Dimensionality reduction

Dimensionality reduction was used to visualize how population activity of subiculum encodes spatial position. t-SNE (45) was applied to the population activity matrix, which was constructed by concatenating population vectors (Equation. (34)) across time. t-SNE was implemented using the sci-kit learn function TSNE. We also used other dimensionality reduction methods for analysis (see supplementary text, figs. S20 and S21).

## Acknowledgments

AB and FW thank Soren Bendt Petersen for help with Blender surface normal wireframe diagrams, and Colin Lever discussion and comments on the manuscript.

## Data and code availability

The code supporting the findings of this study will be made available after publication under https://github.com/Bicanski-NCG. Correspondence should be addressed to AB and FW.

## Appendices

Supplementary Text

Figs. S1 to S21

Tables S1 to S3

References (*90*–*92*)

## Supplementary Text

### A Overview

This supplementary text is structured as follows. In section B, we present extensions of the Disco model. In section C, based on discontinuity detection, we show two alternative/complementary models for subicular cells. In section D, we present additional analyses of simulation results that are not shown in the main text. In section E, we provide the simulation details for the presented results, including environment configurations and simulation durations.

### B Extension of the Disco model

As a general theoretical framework, Disco can be extended to account for a broader range of functions and experimental phenomena. Here, we present three examples: defining neuronal selectivity via discontinuities (i), modifying receptive fields (ii), and altering neuronal firing mechanisms (iii).

#### B.1 Disco corner cells with a broadened notion of discontinuity

As previous work (3) has suggested that a corner arises from the conjunction of two intersecting walls, we restricted corner-cell tuning for horizontal discontinuities to jump discontinuities (*D*_*t,h*_(*θ, φ*)) in the main text. With a finite sampling step size, discrete sampling of surface normals can explain the trace-like representations of inserted corners as the two connected walls gradually separate (Fig. 3G). Alternatively, this effect can also be captured by extending the definition of horizontal discontinuities. Specifically, Equation. (4) is reformulated as

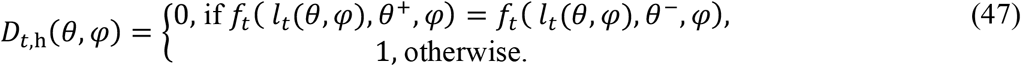

Under this formulation, corner cells respond not only at corner locations, where surface normals from both inserted walls can be sampled, but also in the vicinity of walls, corresponding to transitions from wall to open space, where surface normals are undefined. This model reproduced corner-cell responses to inserted corners (fig. S5) that are more consistent with experimental observations (3).

**Fig. S1.**
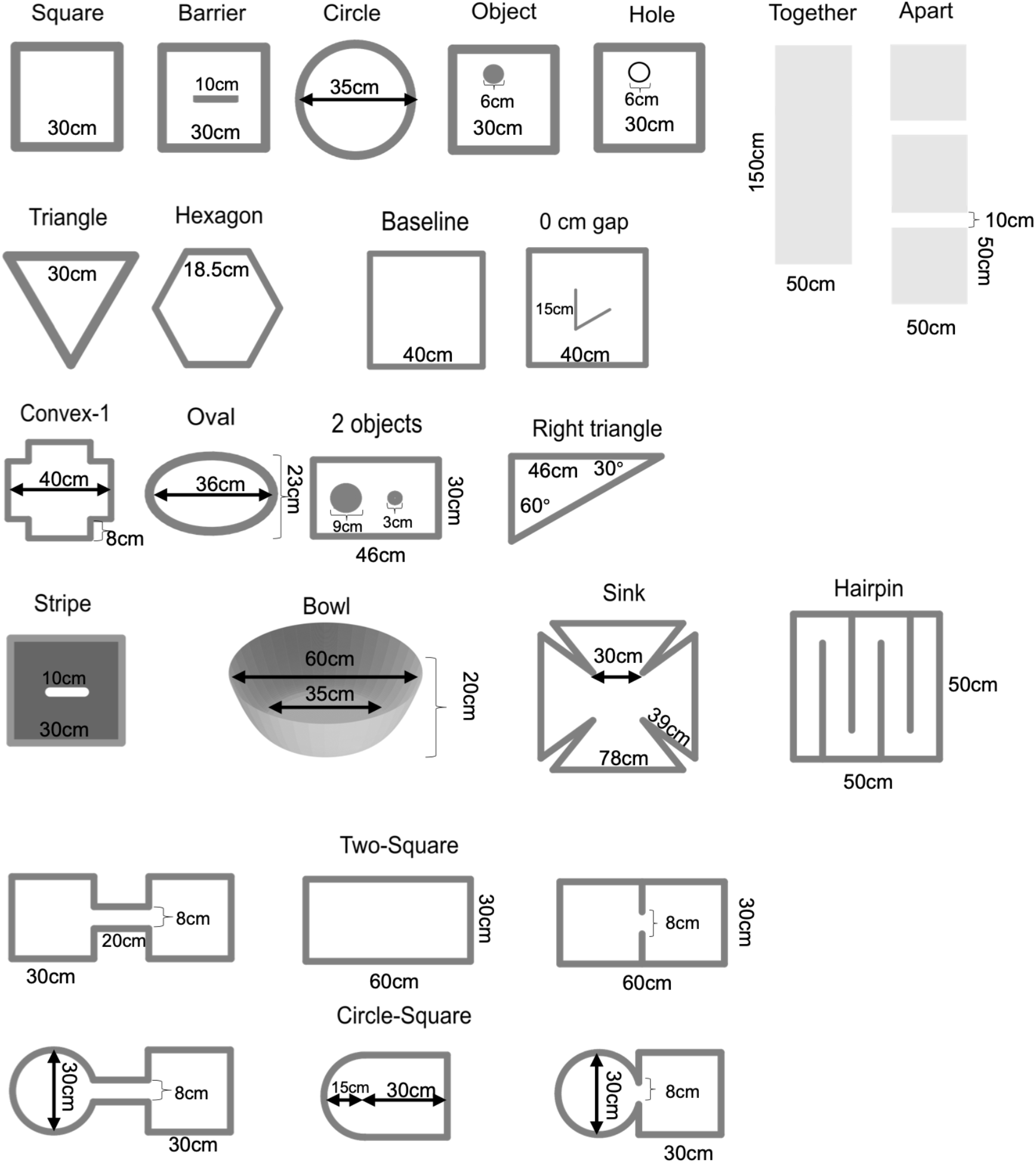
Sizes of the simulated environments used for the majority of the results shown in this study.

**Fig. S2.**
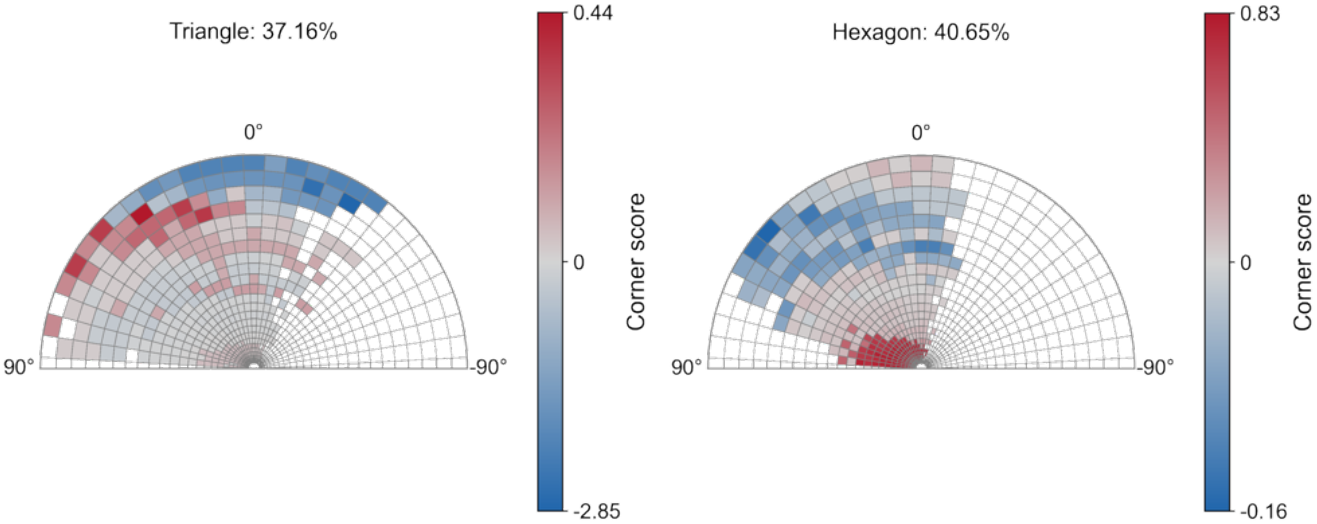
Distribution of corner scores for Disco corner and non-corner neurons in triangular and hexagonal environments. The proportion of corner cells (“corner score” > 0) is indicated above each plot.

**Fig. S3.**
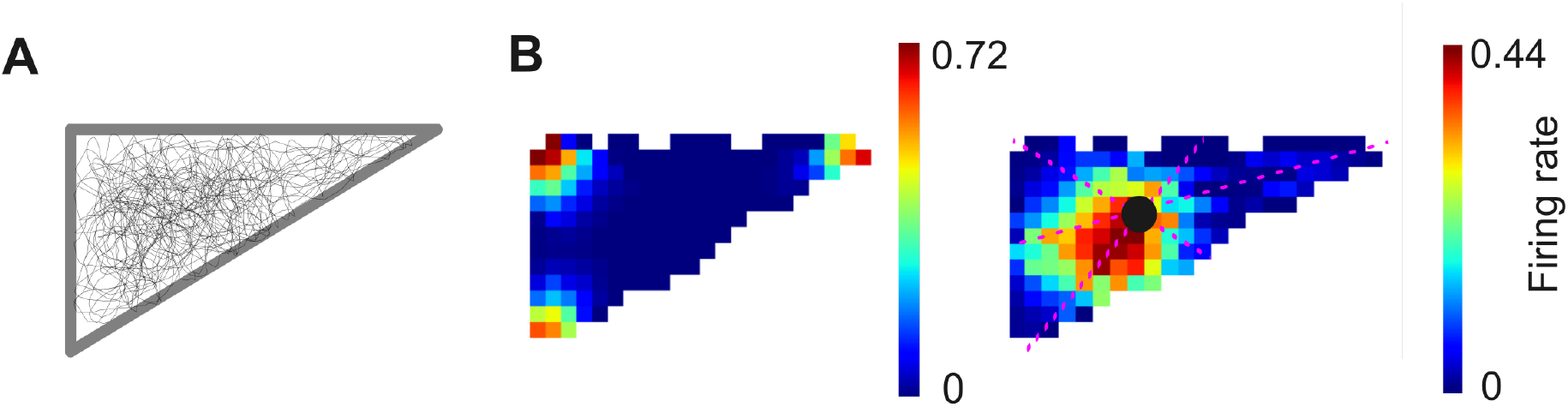
Activities of corner cells in environments with irregular shape. (**A**) Simulated trajectory. (**B**) Left: an example corner cell responding to a discontinuity located at a short distance and positive elevation relative to the agent. Right: an example non-corner cells with positive tuning elevations and long tuning distances. Compared to environments with regular shapes (Fig. 3b), long-range non-corner cells cannot precisely encode the environmental centroid (black dot). Pink dashed lines represent the medians of the triangle.

**Fig. S4.**
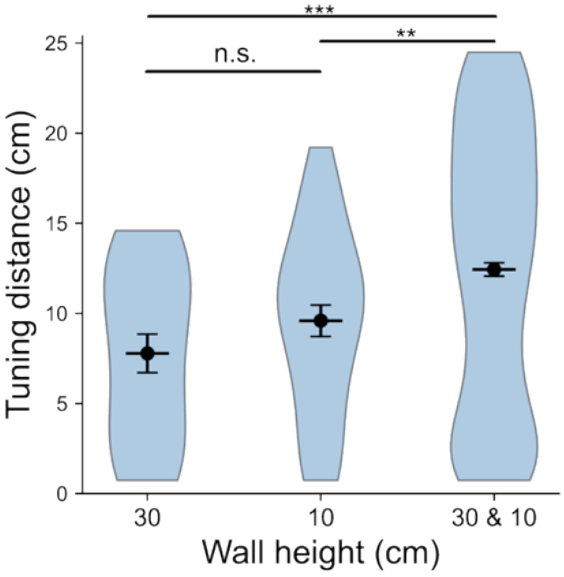
Tuning distances of non-corner cells in square environments with different wall heights. Violin plots depict the density distribution; black dots with error bars indicate mean ± s.e.m. (30cm, n=20 “neurons” ; 10cm, n=33 “neurons” ; 30&10cm, n=439 “neurons”). (30cm versus 10cm: *t*_41.4_ = −1.29, *P* = 0.21, *d* = −0.36; 10cm versus 30&10cm: *t*_44.1_ = −2.96, *P* = 0.00, *d* = −0.43; 30cm versus 30&10cm: *t*_23.6_ = −4.03, *P* = 0.00, *d* = −0.72; 2-sample *t* test) \**P* < 0.05, \*\**P* < 0.01, \*\*\**P* < 0.001.

**Fig. S5.**
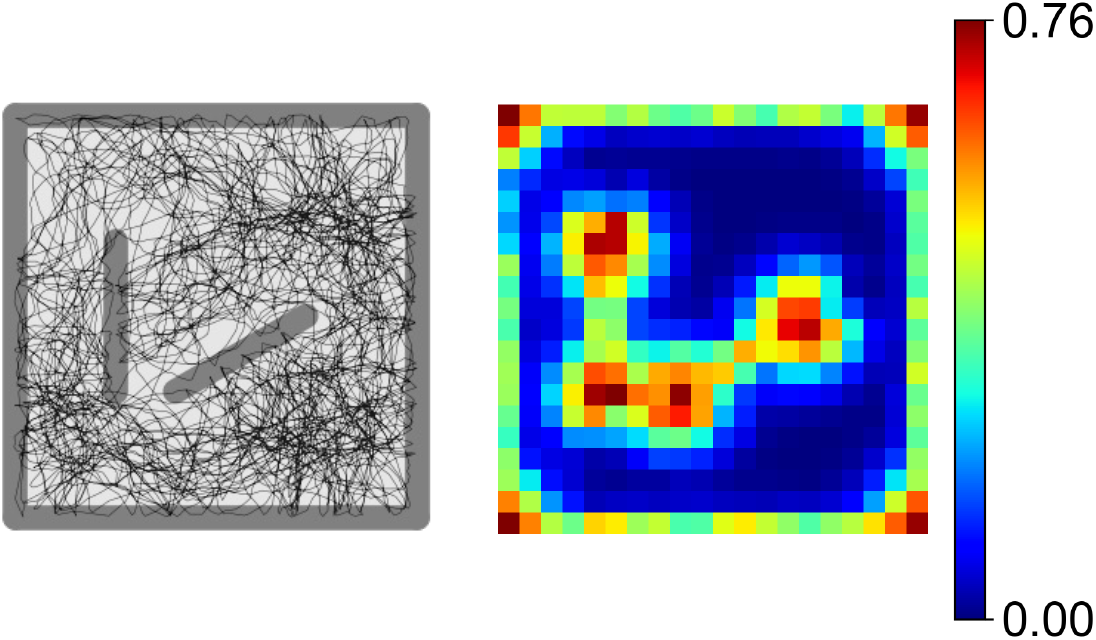
Example Disco corner cell that responds to the corners indicated by the jump discontinuities (edge of barrier to open space). See details in the supplementary text.

#### B.3 Modelling egocentric vertex coding in the Disco framework

Here, we investigate corner coding within an egocentric receptive field. Modeling egocentric boundary coding relies on adjustments of tuning variables, parameterizing the loaction of boundaries in terms of egocentric azimuth and distance (see materials and methods). However, egocentric corner coding in Disco emerges from constraints imposed during the discontinuity detection process. Specifically, we restrict the angular range over which horizontal discontinuities are detected, coupling it to the agent’s allocentric head direction (*HD*_allocentric_) and the neurons’ preferred egocentric bearing (*θ*_*E*_). The horizontal discontinuity contributes to the firing rate of neurons *k* (tuned to distance *r*_*k*_, elevation *φ*_*k*_, egocentric bearing *θ*_*E,k*_), is given by reformulating Eq. (18) as follows:

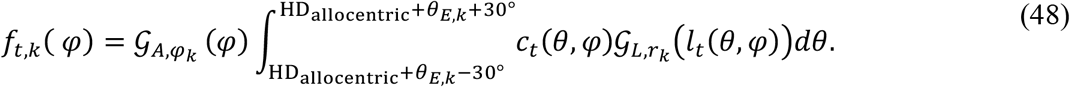

Here, we assume the detection is limited to an angular window of ±30◦. An egocentric bearing of 0◦ corresponds to a discontinuity in front of the agent, and 180◦ denotes one located behind the agent. Bearings of 90◦ and 270◦ represent directions to the left and right of the agent, respectively.

This model variant reproduces the responses of egocentric vertex cells similar to reports from RSC (35) (fig. S6). We did not simulate the effects of concavity/convexity or corner angles, as their firing rates are independent of these factors. Additionally, by using the allocentric receptive field shown in the main text, the model can reproduce the allocentric vertex coding observed in the RSC (35).

#### B.3 Modeling Disco neurons with a threshold-linear transfer function

We extend the tuning functions of Disco BVCs and (non-) corner cells (Equations. (10) and (19)) to further modulate neuronal responses. The firing rates of the simulated neurons are determined by a threshold-linear transfer function:

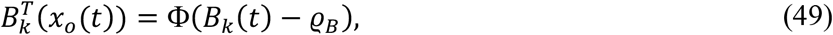

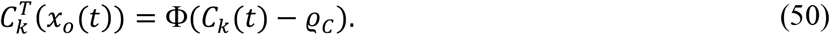

Here,ϱ_*B*_, ϱ_*C*_ and Φ(·) provide a threshold-linear transfer function.Φ(·) is defined as

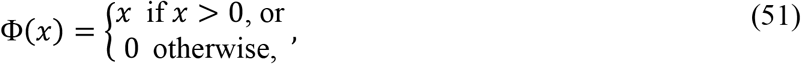

where ϱ_*B*_ and ϱ_*C*_ denote the threshold constants for Disco BVCs and (non-)corner cells, respectively. When the thresholds are set to zero, the model reduces to the formulation described in the main text.

To examine the functional role of this thresholding mechanism, we applied this extended model to our simulation in the natural environment (Fig. 4). We found that the threshold values (ϱ_*B*_, ϱ_*C*_) control the resolution of geometric encoding (figs. S13-S14). In environments with noisy geometric information, increasing the threshold emphasizes the global geometric structure of the environment while suppressing fine-scale details. This indicates that thresholding mediates a trade-off between encoding fine details and capturing coarse, global geometric features.

### C Alternative ways for modelling subicular cells

Here, we first show that corner cells can be modelled as neurons with a ring-like receptive fields. Then we present an alternative implementation for modeling BVCs based on the Disco theory. Moreover, inspired by BVC–place cell (PC) models (17,38,90) and BVC–vertex cell model (35), in which the firing rate of a PC/vertex cell) is modeled as a thresholded sum of the firing rates of connected BVCs, we construct a network model within the same framework. Specifically, we model other subicular cells (SCs) as the weighted sums of the firing rates of Disco BVCs. We found that this model can account for the emergence of corner cells, with firing rates that depend on wall height. Unless otherwise specified, most model parameters are consistent with those in the main text (table S1), and newly introduced parameters are listed in table S3.

**Table S1.** Model parameters.

| Parameter | Description | Default |
| --- | --- | --- |
| $L$ | Upper bound on detection vector length | 0.3m |
| $dl$ | Numerical step size of detection vector length | 0.01m |
| $d\theta$ | Numerical azimuthal integration step | $10^\circ$ |
| $d\varphi$ | Numerical elevation integration step | $2^\circ$ |
| $\xi$ | Inverse gradient in the linear function of tuning distance of neurons | 10 |
| $\sigma_0$ | Intercept in the linear function of tuning distance of neurons | 0.03m |
| $d\theta^h$ | Numerical head-centered azimuthal integration step | $10^\circ$ |
| $dt$ | Numerical time step | 0.1s |
| $N_A$ | Numbers of Disco axis cells contributed to event boundary detection | 600 |
| $N_B$ | Numbers of Disco BVCs contributed to event boundary detection | 600 |
| $N_C$ | Numbers of Disco corner and non-corner cells contributed to event boundary detection | 900 |
| $w_A$ | Scaling factor for Disco axis cell contributions to event boundary detection | 0 |
| $w_B$ | Scaling factor for Disco BVC contributions to event boundary detection | 1 |
| $w_C$ | Scaling factor for Disco corner and non-corner cell contributions to event boundary detection | 10 |

**Table S2.** Simulation durations for the majority of shown results.

| Environment | Simulation duration | Referenced figure(s) |
| --- | --- | --- |
| Square | 10min | Figs. 2B, M-L, 3A; figs. S7, S9-S11 |
| Barrier | 10min | Fig. 2B |
| Circle | 10min | Fig. 2B |
| Object | 10min | Fig. 2B |
| Hole | 10min | Fig. 2E |
| Together | 30min | Fig. 2C, G-H |
| Apart | 30min | Fig. 2D |
| Triangle | 5min | Fig. 3A |
| Hexagon | 10min | Fig. 3A |
| Baseline | 15min | Fig. 3G |
| 0/1.5/6 cm gap | 15min | Fig. 3G, fig. S5 |
| Convex-1 | 10min | Fig. 3I, fig. S6 |
| Oval | 10min | Fig. 3K |
| 2 objects | 10min | Fig. 3M |
| Right triangle | 5min | fig. S3 |
| Stripe | 10min | Fig. 5B |
| Bowl | 10min | Fig. 4B-C |
| Sink | 20min | Fig. 4F-H |
| Two-Compartment environments | 20min | Fig. 6A-G; figs. S18-S21 |

**Table S3.** Additional parameters introduced in alternative models of subicular cells.

| Parameter | Description | Default |
| --- | --- | --- |
| $L_b$ | Length of surface-travelling rays | 1m |
| $dl_b$ | Numerical distance integration step | 0.01m |
| $d\theta_b$ | Numerical azimuthal integration step | 6° |
| $N_{\text{BVC}}$ | Numbers of BVCs | 720 |
| $N_{\text{SC}}$ | Numbers of SCs | 300 |
| $n_b$ | Number of BVCs connected to each SC | 10 |
| $T_b$ | threshold | 1.3 |

#### C.1 Modelling corner cell with the ring-like receptive fields

Since we allow for variations among different Disco cell types (assuming they sample different kinds of discontinuities), we also test modifications of classical BVC selectivity, to see if it can be adapted to produce other cell types. Instead of assuming that corner cells are selective to the elevation and distance of discontinuities detected by the horizontal discontinuity indicators *D*_*t,h*_(*θ, φ*), an alternative approach for generating corner representations is to use classical BVC-like receptive fields without angular selectivity (here, augmented by discontinuity detectors).

In this model, we assume that neurons respond to discontinuities determined by the vertical discontinuity indicators *D*_*t,v*_(*θ, φ*), which are also used by BVCs for boundary detection (materials and methods). Given a ring-like receptive field, these cells respond to discontinuities detected in any direction. The tuning function includes only one Gaussian component that responds to the vector distance (*l*_*t*_(*θ, φ*)). The vertical discontinuity contributes to the firing rate of a unit *k* (tuned to distance *d*_*k*_, azimuth *ϑ*_*k*_) as follows.

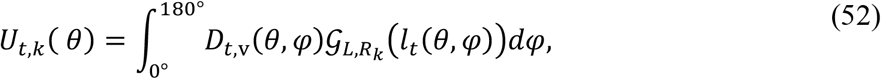

In our simulation (fig. S6), the tuning distances *R*_*k*_ increased linearly from 0.95 cm to 24.97 cm over a radial extent of 22 distance units. At time *t*, the contribution of all detected discontinuities to the firing of the unit *k* is determined by integrating Equation. (51) over *θ*:

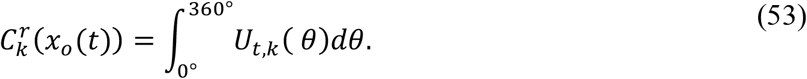

**Fig. S6.**
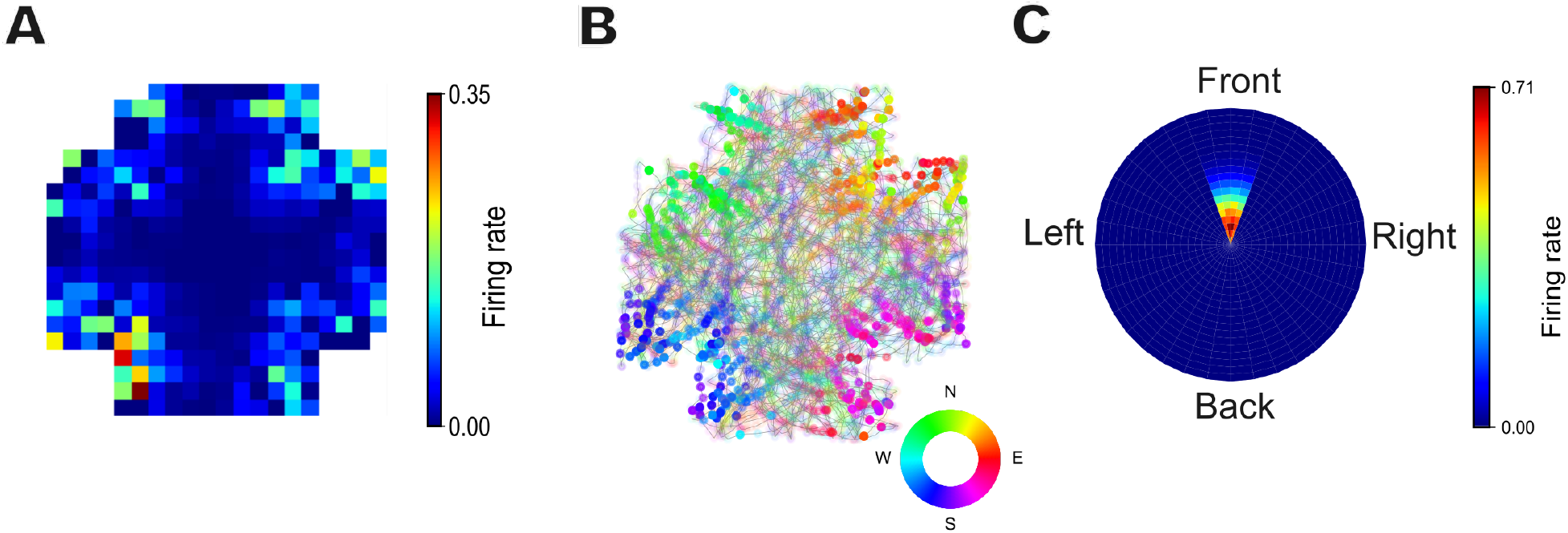
Example egocentric vertex cell in the Disco framework. (A) Allocentric rate maps. (B) Trajectory plots with neuronal firing color-coded by head direction, with transparency proportional to firing rate. (C) Egocentric vertex rate maps, where firing rate is represented as a function of proximal vertex locations in polar coordinates relative to the agent. The example neuron has a preferred egocentric bearing of 0° (see details in the supplementary text).

Here, neurons with short tuning distances show corner representations (fig. S7), as the agent detects more discontinuities near corners (i.e., at the intersection of two walls) than along a single straight wall. Corner cells appear as a variant of BVCs under this formulation. Thus, classical BVCs with less precise angular selectivity (in the extreme case, 360° selectivity at a given distance) can give rise to corner cell responses after thresholding. However, this model cannot detect the surface normals of the two walls forming a corner and therefore fails to account for the effects of wall height, corner angle, and the non-overlapping coding of convex and concave corners. Specifically, regarding the latter point, such a neuron would exhibit a firing field on both sides of an inserted corner.

**Fig. S7.**
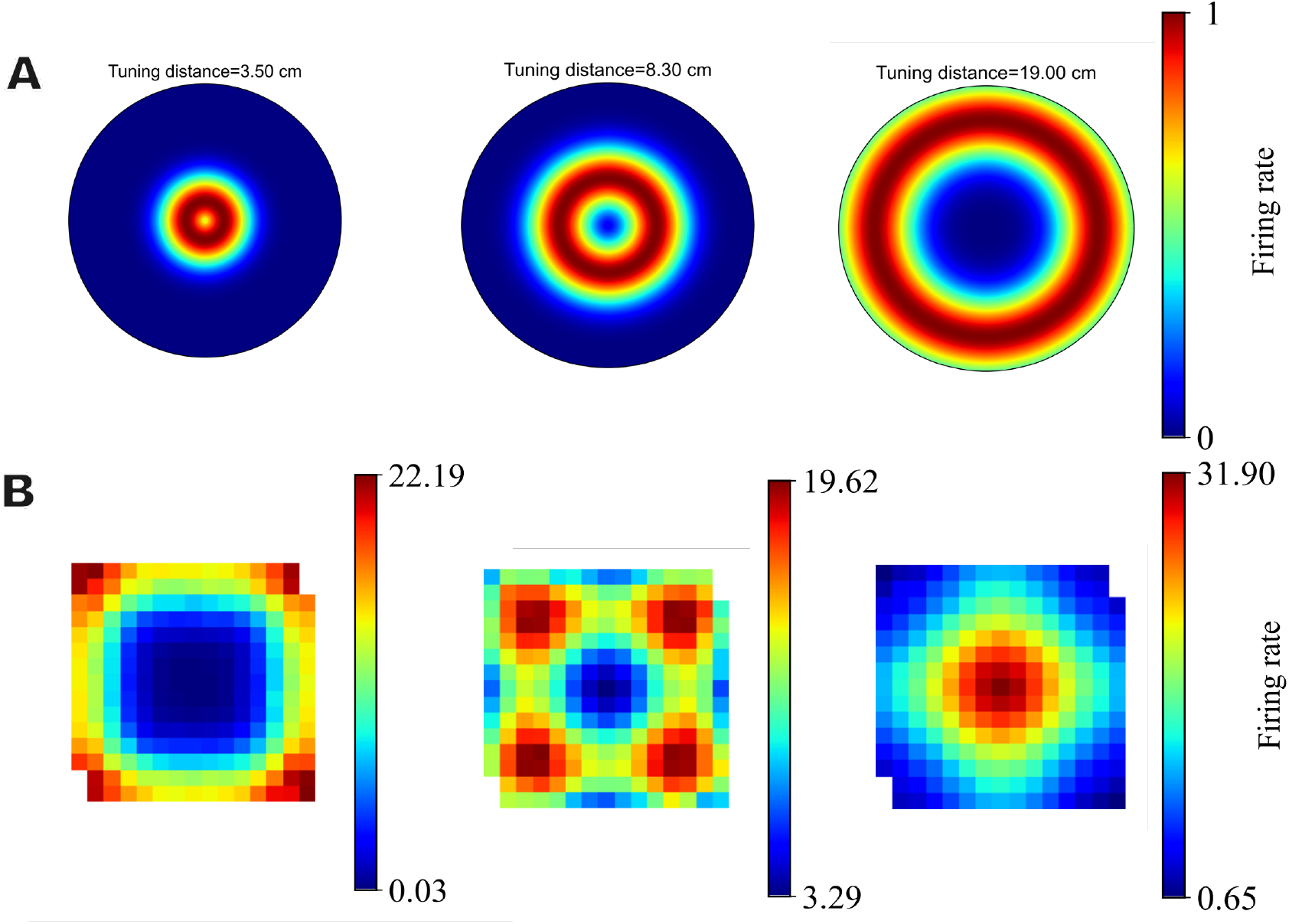
Corner cells generated from the ring-like receptive fields. (**A**) Example neurons with receptive fields of different tuning distances (see details in supplementary text). (**B**) Rate maps. We simulated an agent exploring a square environment. Neurons with short tuning distances show corner representations (left), whereas neurons with longer tuning distances can encode other geometric features, for example the center of a square environment.

#### C.2 An alternative way for modelling BVCs

In this model variant, instead of using the detection vectors defined in the spherical coordinate system (*l, θ, φ*) to sample the surface normals (materials and methods), we assume that BVCs detect discontinuities in a quasi-planar manner. Specifically, we define surface-travelling rays of length *L*_*b*_ emitted from the agent’s body centroid *x*_*o*_(*t*), each characterized by a fixed azimuthal direction *θ*. These rays propagate along the local tangent direction of the perceptual surfaces. When a ray encounters surfaces with varying surface normals (e.g., transitions from the floor to the walls), it continues to propagate in the same azimuthal direction (fig. S8).

**Fig. S8.**
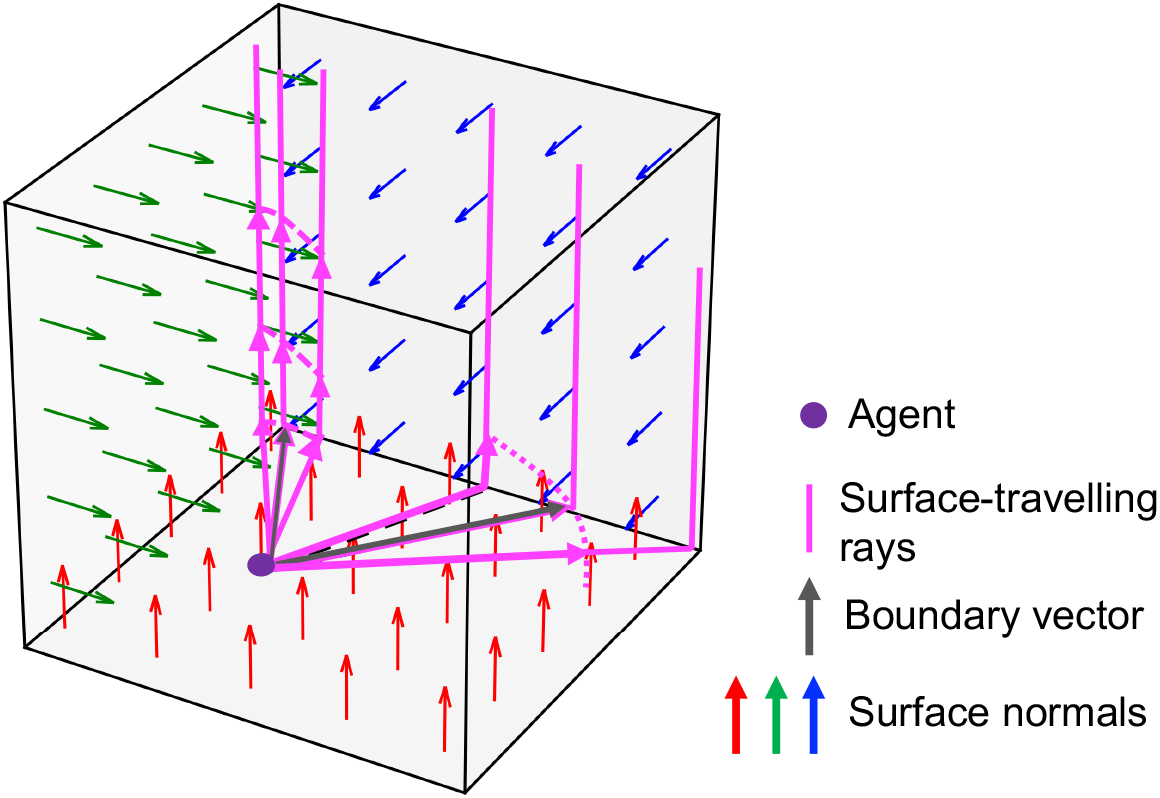
Model schematic of the alternative BVC model. BVCs respond to discontinuities detected by comparing adjacent surface normals, which are sampled by equal-length vectors (pink arrows) defined along surface-travelling rays (see supplementary text for details). Unlike the BVC model described in the main text (materials and methods), multiple discontinuities can be detected along a fixed azimuthal direction (e.g., at environmental corners). In the BVC–SC network model, the number of detected discontinuities modulates synaptic weights and facilitates the emergence of corner cells (see supplementary text for details).

With this sampling scheme, all sampled points lie on the surfaces of the environment (e.g., walls or floor). Along a given ray, each point *x*_*t*_(*θ, l*), with *θ* ∈ [0, 360^◦^] and *l* ∈ [0, *L*_*b*_], is associated with a feature value 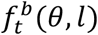, as defined in the materials and methods. For our simulations in spatial domains, 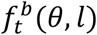 represents the surface normal.

We defined the discontinuity indicator that determines the BVCs’ firing rates as follows,

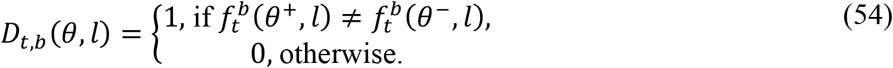

For instance, in a square environment, if there exists a point *x*_*t*_(*θ, l*) such that *D*_*t,b*_(*θ, l*) = 1 and *v*_*t*_(*θ, l*) = (0,0,1), then a boundary (corresponding to the “floor” panel) is present in the allocentric direction *θ* at distance *l*. As shown in fig. S8, multiple discontinuities may occur in corner regions. Here, consistent with the previous models, we assume that only the boundary closest to the agent contributes to the firing rate of BVCs for direction *θ*, with the corresponding distance denoted by *l*_*t*_(*θ*).

Then, the contribution of the detected discontinuity to the firing rate of the BVC *k, B*_*k*_ (Eq. (6)), is reformulated as:

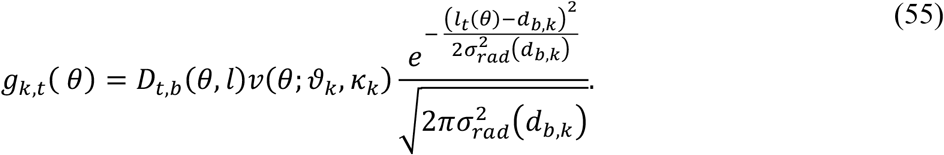

Here, the tuning azimuths were uniformly distributed with a spacing of 6◦,*ϑ*_*k*_ ∼ 𝒰[0◦, 360◦]. The tuning distances (*d*_*b,k*_) increased linearly from 2.64 cm to 23 cm over a radial extent of 12 distance units. This model variant reproduces BVCs’ boundary representations for walls (fig. S8), and can also be applied to model the selectivity to objects, drops and stripes.

#### C.3 The BVC-SC network model

We build the BVC-SC network based on the BVC-PC model. In our simulation, the subiculum consists of *N*_*BVC*_ BVCs and *N*_*SC*_ SCs, and each SC receives inputs from a selection of *n*_*b*_ BVCs. Following previous model, the firing of SC unit *j* at location *x*_*o*_(*t*) (referred to as *F*_*j*_(*t*)) was derived as the thresholded sum of the connected BVCs,

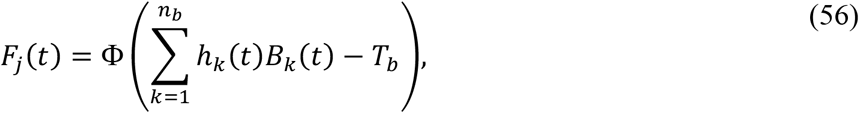

where threshold *T*_*b*_ is costant, and Φ is the activation function defined as

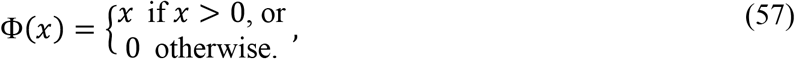

BVC–vertex cell model can reproduce the corner representation. However, unlike vertex cells, the firing rates of corner cells are affected by more geometric features (e.g., wall height). In this model variant, Disco provides a measure of wall height. More discontinuities are present at a corner compared to the wall–floor intersection (fig. S8). To capture the effect of wall height on corner cell activity (3), we introduce a weighting parameter *h*_*k*_(*t*) to describe the contribution of BVC *k* to SCs, which is given by:

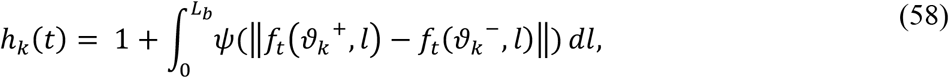

where

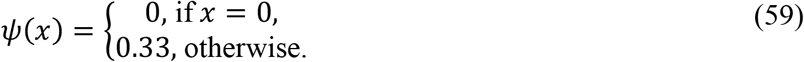

If *h*_*k*_(*t*) = 1, this indicates that no discontinuity is present in the direction *ϑ*_*k*_. If *h*_*k*_(*t*) = 1.33, a single discontinuity is present (e.g., the intersection between a wall and floor). Otherwise, multiple discontinuities are present in this direction (e.g., a corner). Corners composed of higher walls correspond to larger values of *h*_*k*_(*t*), leading to stronger firing of the connected SCs.

#### C.4 Corner cells, place cells and BVCs can emerge from the network model

The BVC-SC model can reproduce PC activity patterns (fig. S9) consistent with previous studies (17,90). In addition, we found a subset of SCs exhibiting diffuse place fields, consistent with experimental findings that subiculum PCs exhibit multiple spatial fields compared to hippocampal PCs (73). Moreover, we found SCs that exhibits corner-cell-like and BVC-like activity (figs. S10 and S11).

**Fig. S9.**
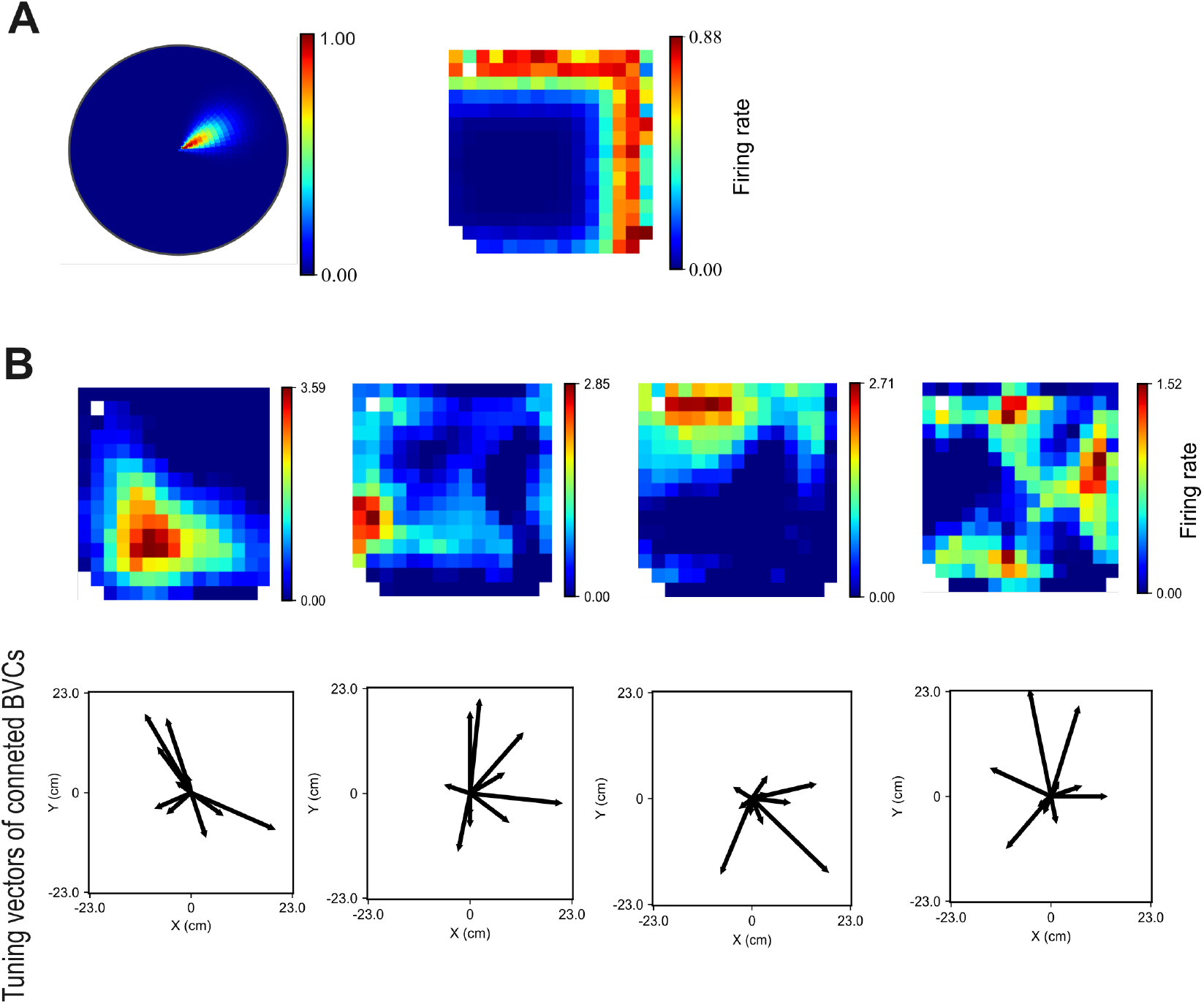
Place cells generated from the thresholded sum of BVCs. (**A**) Receptive field and firing rate map of an example BVC (supplementary text). (**B**) Top: Rate maps of example place cells emerging from the BVC-SC network model (supplementary text). Bottom: Vector plots indicate distance and directional selectivity of randomly chosen input BVCs that feed into the given place cell.

**Fig. S10.**
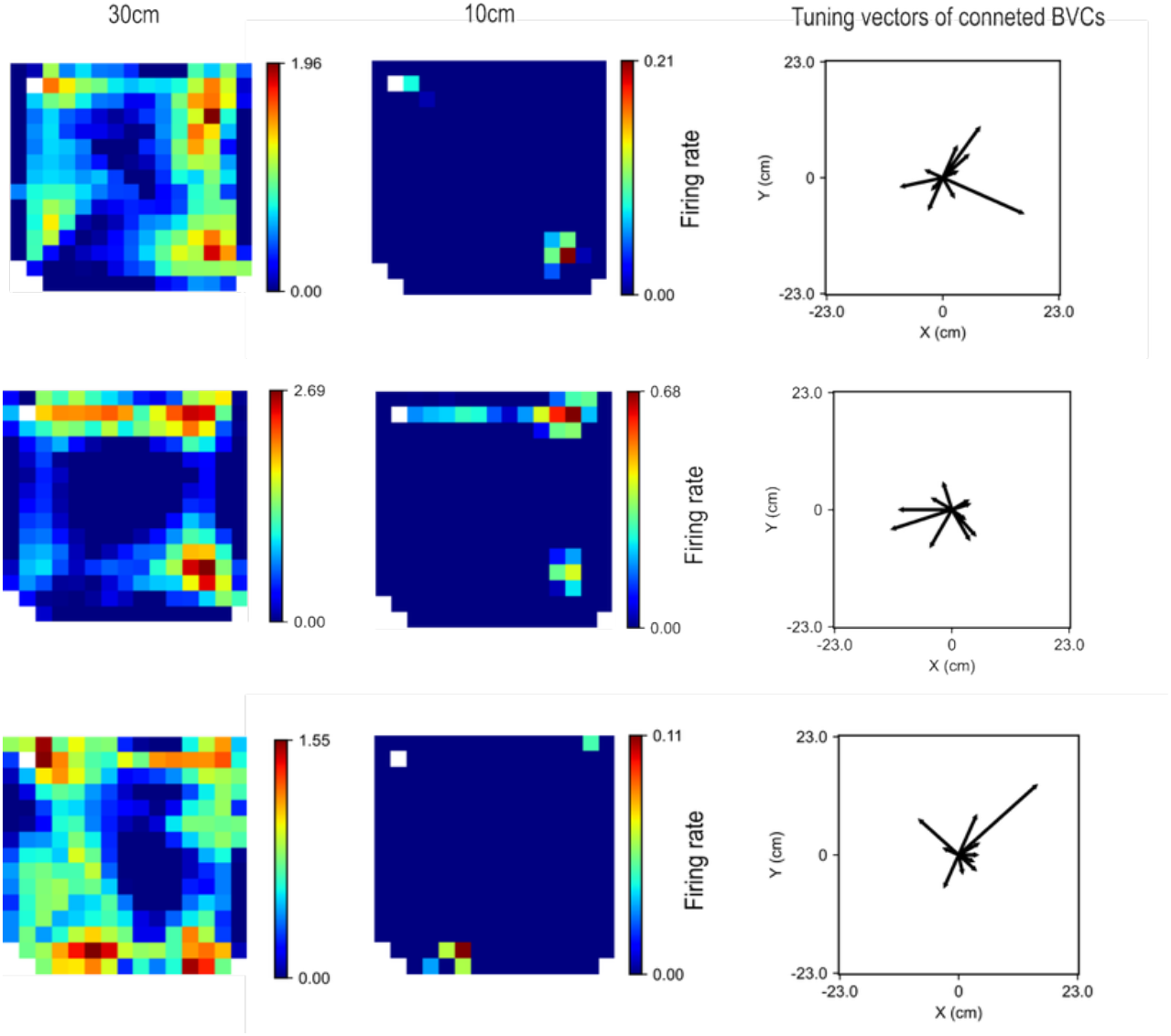
Alternative corner cell model: corner cells emerging from the BVC-SC network model. Three example neurons exhibited corner representations in the high-wall environment (30 cm; left), which decayed in the low-wall environment (10 cm; middle). Vector plots (right) indicate distance and directional selectivity of randomly chosen input BVCs that feed into the given cell.

**Fig. S11.**
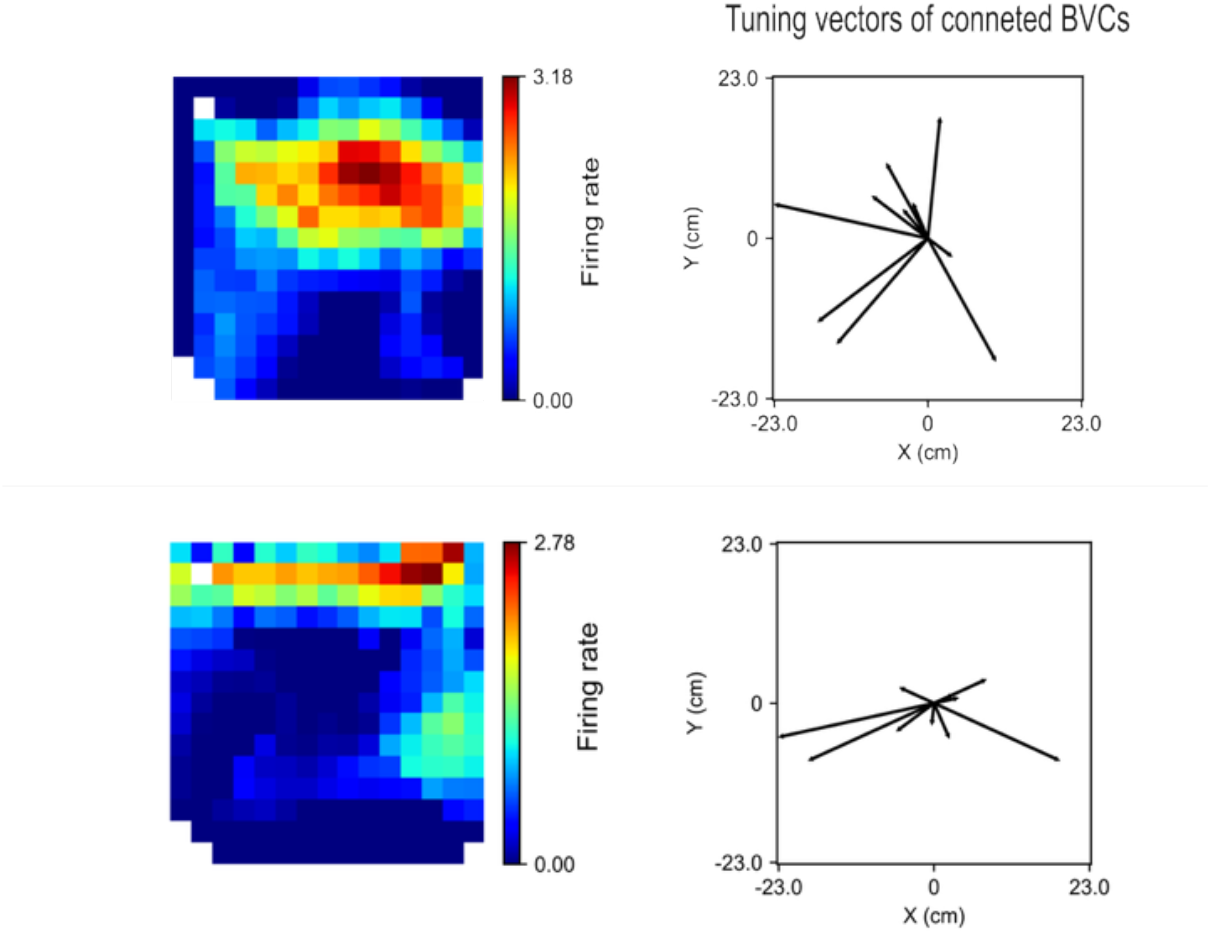
Boundary representations emerging from the BVC-SC network model. Left: Rate maps of example neurons that generate BVC-like activities. Right: Vector plots indicate distance and directional selectivity of randomly chosen input BVCs that feed into the given neuron.

**Fig. S12.**
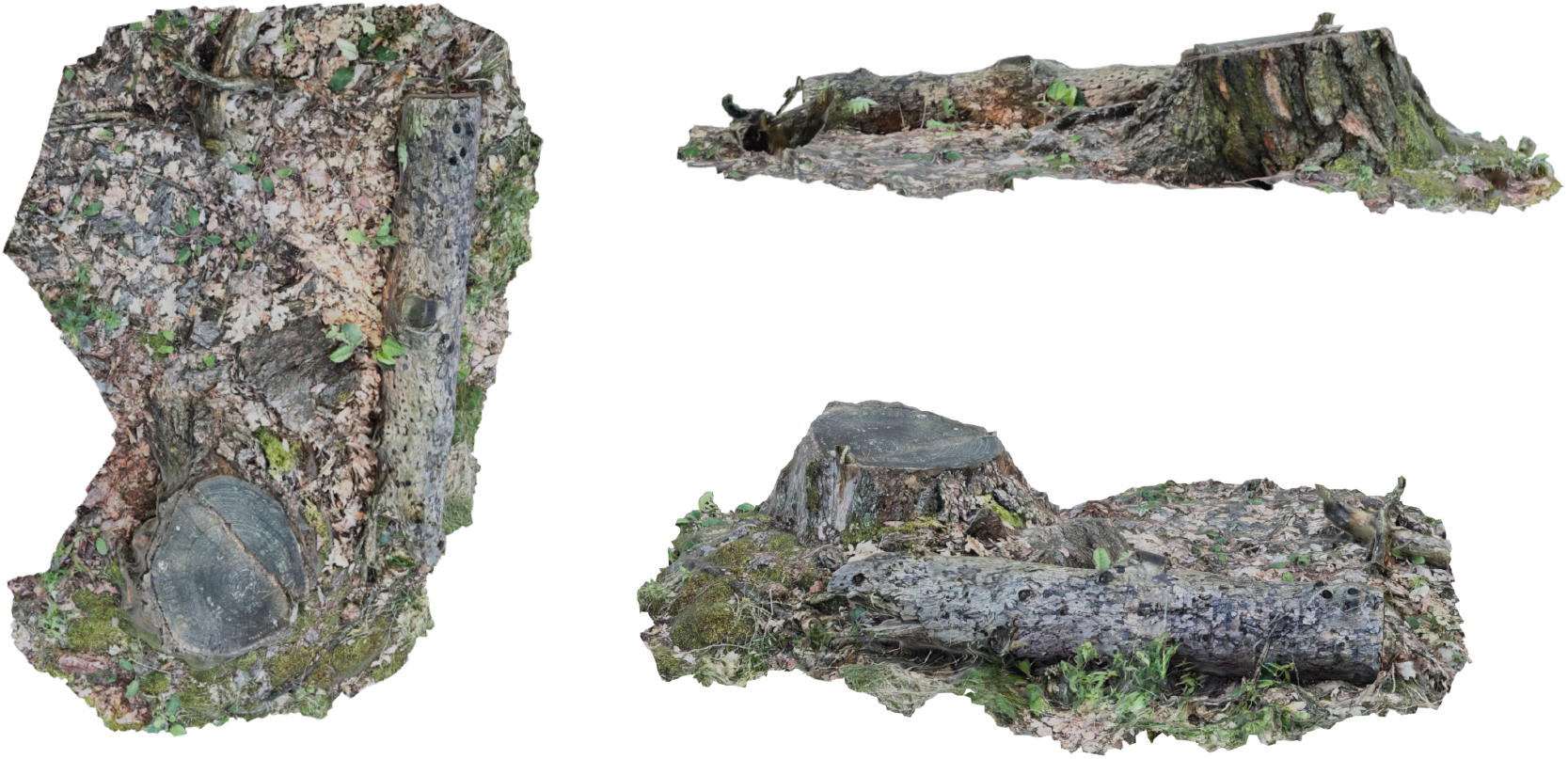
Details of the used nature environment material. Free asset from BlenderKit (https://www.blenderkit.com) under a Royalty Free license.

**Fig. S13.**
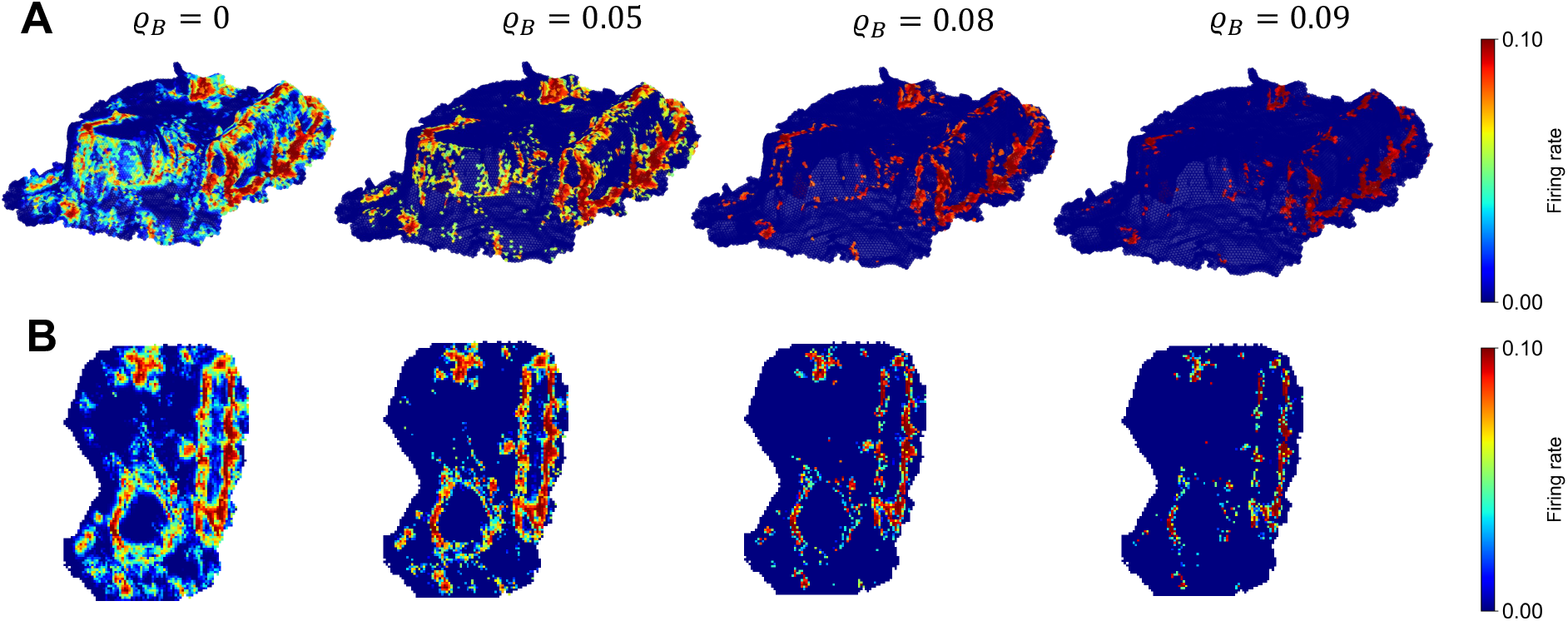
Noise reduction in natural environments by thresholding - BVCs. Assuming a variable input gain (here implemented as thresholding ϱ_*Y*_, see supplementary text), activities of BVCs can facilitate geometric coding with different resolution in natural environments. (**A**) 3D rate map. (**B**) Bird’s-eye-view rate map.

**Fig. S14.**
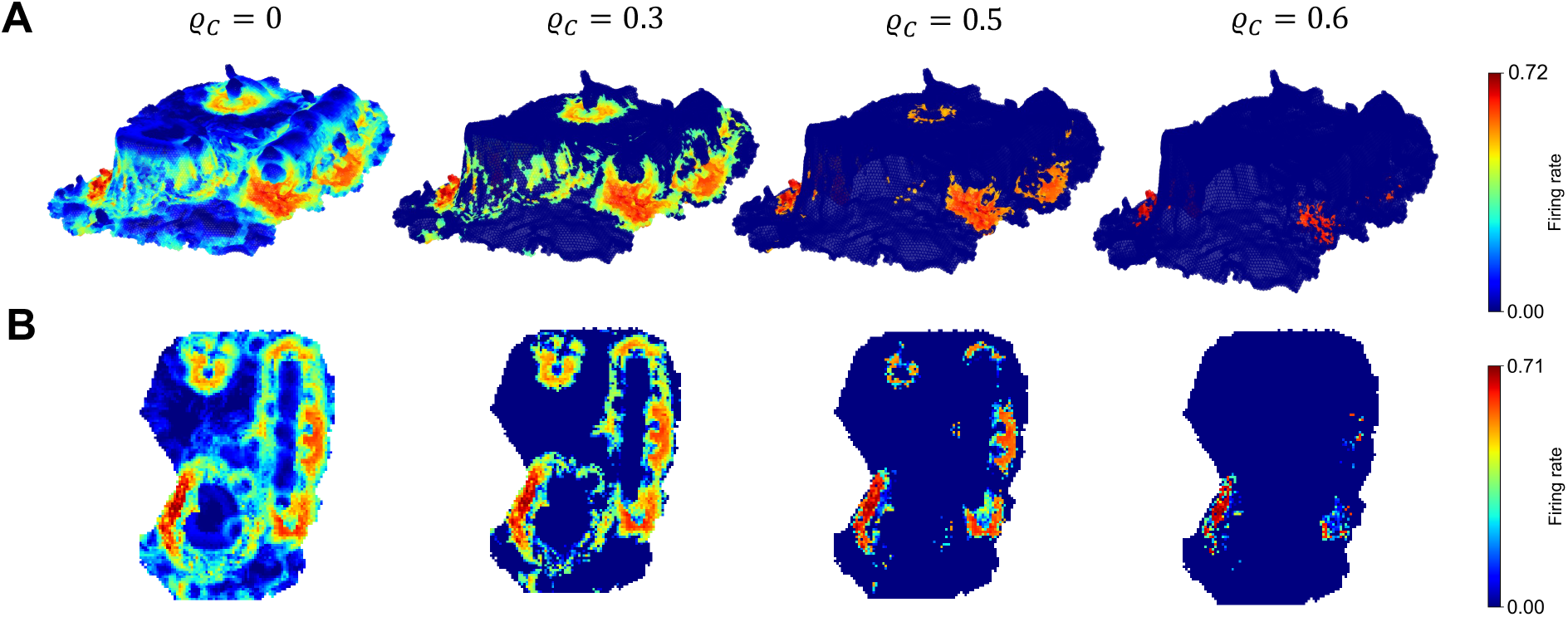
Noise reduction in natural environments by thresholding – corner cells. Assuming a variable input gain (here implemented as thresholding ϱ_*C*_, see supplementary text), activities of corner cells can facilitate geometric coding with different resolution in natural environments. (**A**) 3D rate map. (**B**) Bird’s-eye-view rate map.

By examining their network connectivity, we observe that, compared with BVC-like and PC-like neurons, corner-cell-like SCs preferentially receive inputs from Disco BVCs with shorter tuning distances and tuning directions aligned toward corners. This results in the activation of these neurons when the agent is near any environmental corner. We only simulated exploration trajectories in a square environment. Having introduced a weighting parameter related to wall height (*h*_*k*_(*t*)) for the alternative model, we find that corner representations observed in some SCs in high-wall (30 cm) environments disappear in low-wall (10 cm) environments (fig. S10). This is consistent with experimental observations showing a higher proportion of subicular corner cells in high-wall environments (3).

These results show potential (albeit limited, see below) for complementary ways to generate, for example, corner cells within subiculum through local interactions. Potentially, this mechanism could work alongside neural selectivity for discontinuities. However, relying exclusively on fixed synaptic connectivity may not guarantee corner representations across different environments, would require plasticity in response to changes in environmental geometry, and seems counter to the regularity of corner responses across different shapes, their rapid emergence, as well as their curvature-dependent responses to continuous surfaces. Future work must investigate recurrent network structures in the subiculum to further elucidate the interactions among BVCs, corner cells, and PCs.

### D Additional analysis

#### D.1 Validation of the model using experimental Data

We analyzed data from one animal for the M-maze task (44), two animals for the wagon-wheel task (44), and two animals for the open-field task (43). Results from one representative animal are shown in Fig. 5 and fig. S16, with results from the other animals shown in fig. S15. All trajectory datasets used here reproduce Disco axis cells, as the animals exhibited non-zero acceleration during movement.

**Fig. S15.**
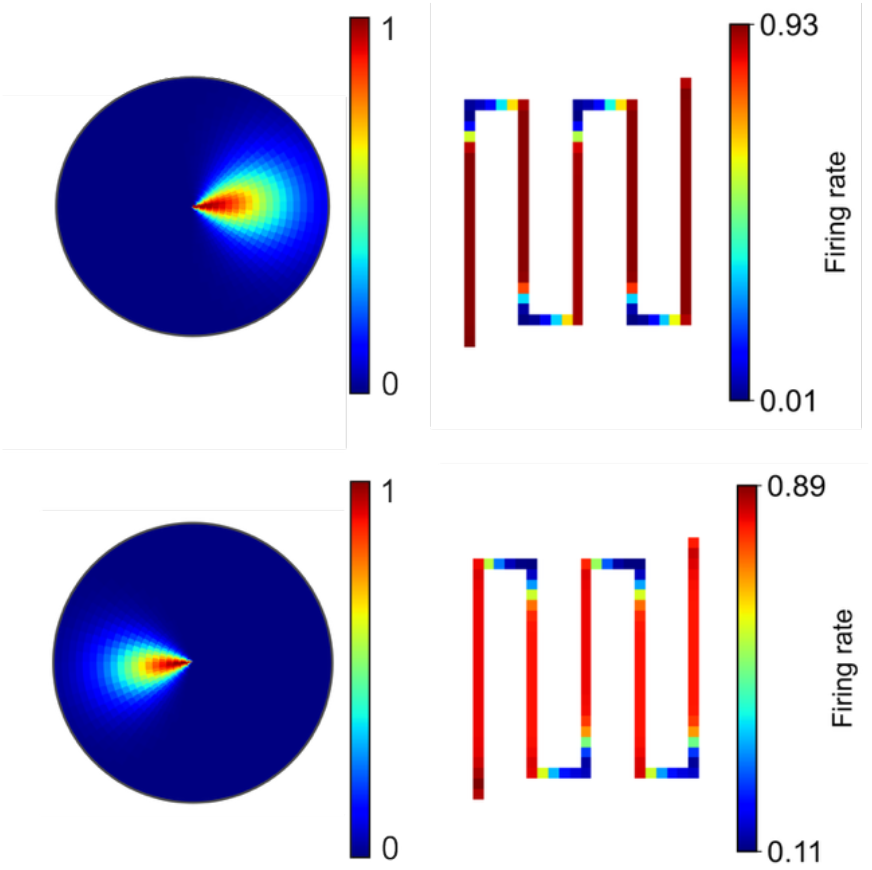
Axis coding by short-range BVCs. A subset of BVCs with short tuning distance can perform axis coding during exploration the linear track. Unlike Disco axis cells (Fig. 5), these neurons also exhibit regular boundary representations in open-field environments. Consistent with this, a previous experimental study (19) showed that a subset of axis-tuned neurons also display BVC-like activity in open fields.

**Fig. S16.**
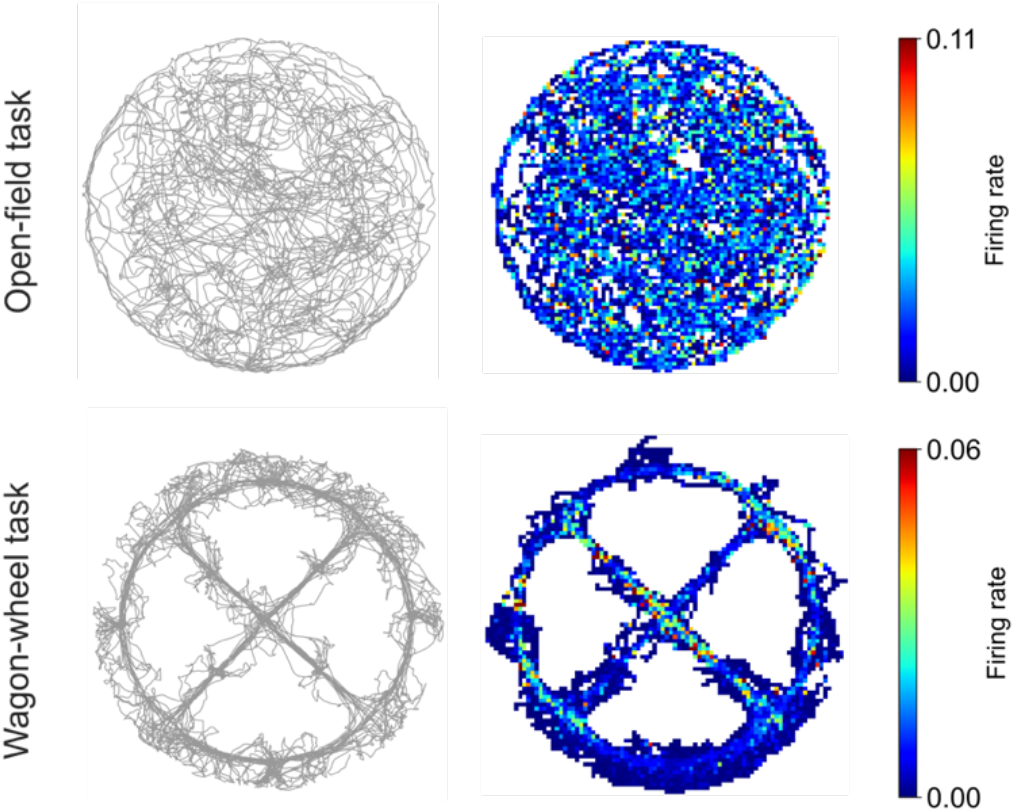
Verification of Disco axis cells by additional experimental trajectory data. Left: Trajectories. Right: Rate maps of simulated Disco axis cells fitted to experimental trajectory data (not shown in the main text) for wagon-wheel(44) and open-field(43) tasks.

#### D.2 Analysis of neural populational activity

We applied dimensionality reduction methods to visualize how population activity in the subiculum encodes spatial position. In addition to t-SNE (materials and methods), we also used two alternative approaches: principal component analysis (91) (PCA) and uniform manifold approximation and projection (92) (UMAP). UMAP was implemented using the umap-learn function *UMAP*, and PCA was implemented using the scikit-learn function *PCA*.

We found that both PCA and UMAP can capture certain geometric features of the two-compartment environment (figs. S20 and S21). Specifically, two identical compartments exhibited overlapped representations, whereas compartments with different shapes produced clearly separable representations. However, compared with two other nonlinear dimensionality reduction methods, PCA failed to separate circular and square arenas when objects were inserted. Moreover, in the two-square environments (Corridor/Adjacent), the representations of the two squares were nearly indistinguishable, indicating that PCA failed to extract allocentric coordinate information distinguishing the two compartments. These results imply the nonlinear characteristics of subicular population activity.

**Fig. S17.**
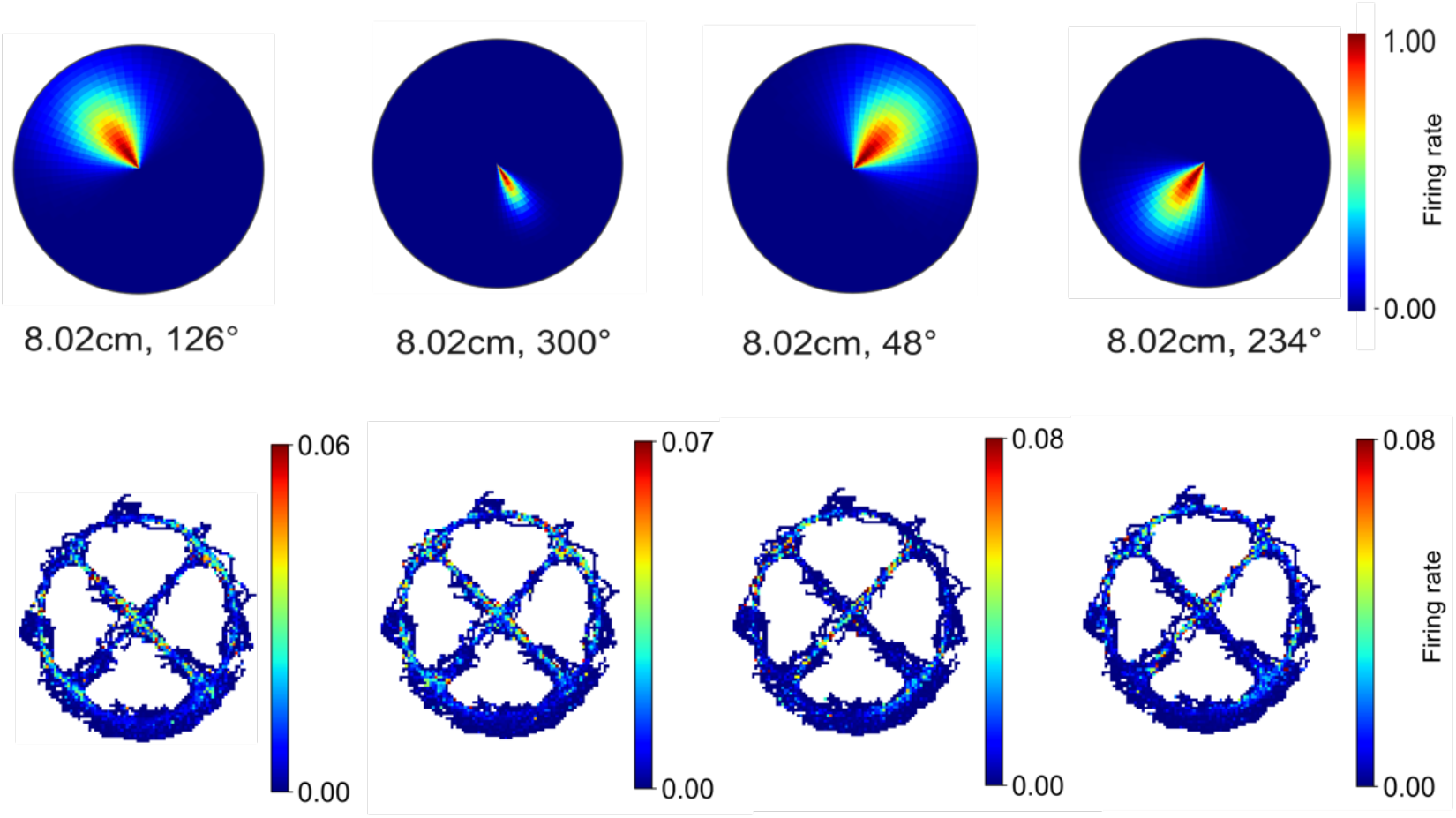
Axis coding on the wagon wheel maze. Vector plots (Top) and rate maps (Bottom) of example axis cells fitted to the Wagon-wheel task trajectory data(44) shown in Fig. 5h. The magnitude and direction of the preferred acceleration-tuning vector are indicated below the vector plots.

**Fig. S18.**
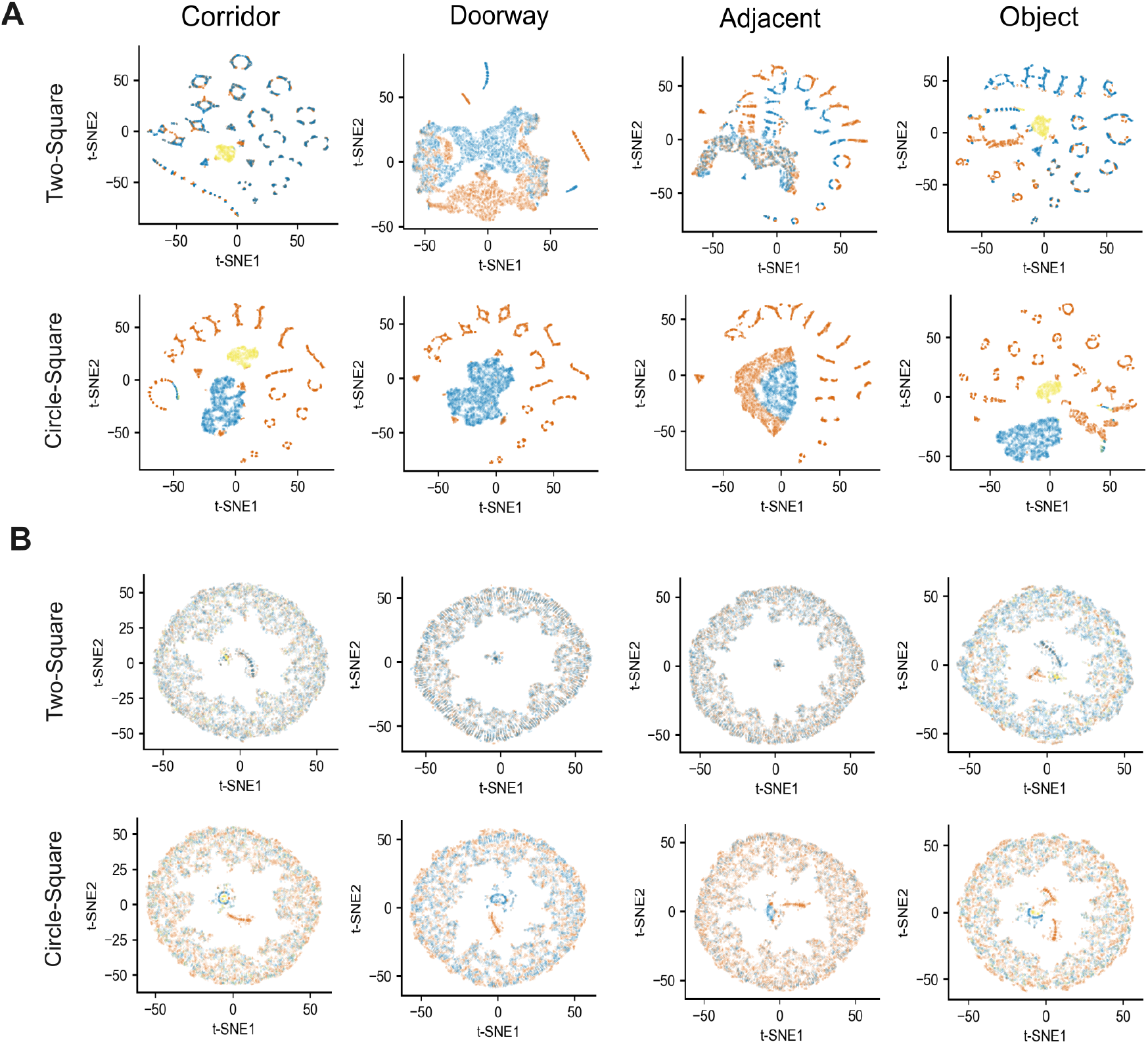
T-SNE dimensionality reduction with the effects of Disco axis cells. Two-dimensional embedding (t-SNE) of population activity in the two-compartment arenas including the influence of Disco axis cells. Each dot represents the population state at a single position. Arena regions of agent positions are color-coded as Fig. 6. t-SNE was conducted on the proposed a population vector, M(*t*) (materials and methods). (**A-B**) Scaling factor of axis cells was set to *w*_*A*_ = 1 (A) and *w*_*A*_ = 100 (B).

**Fig. S19.**
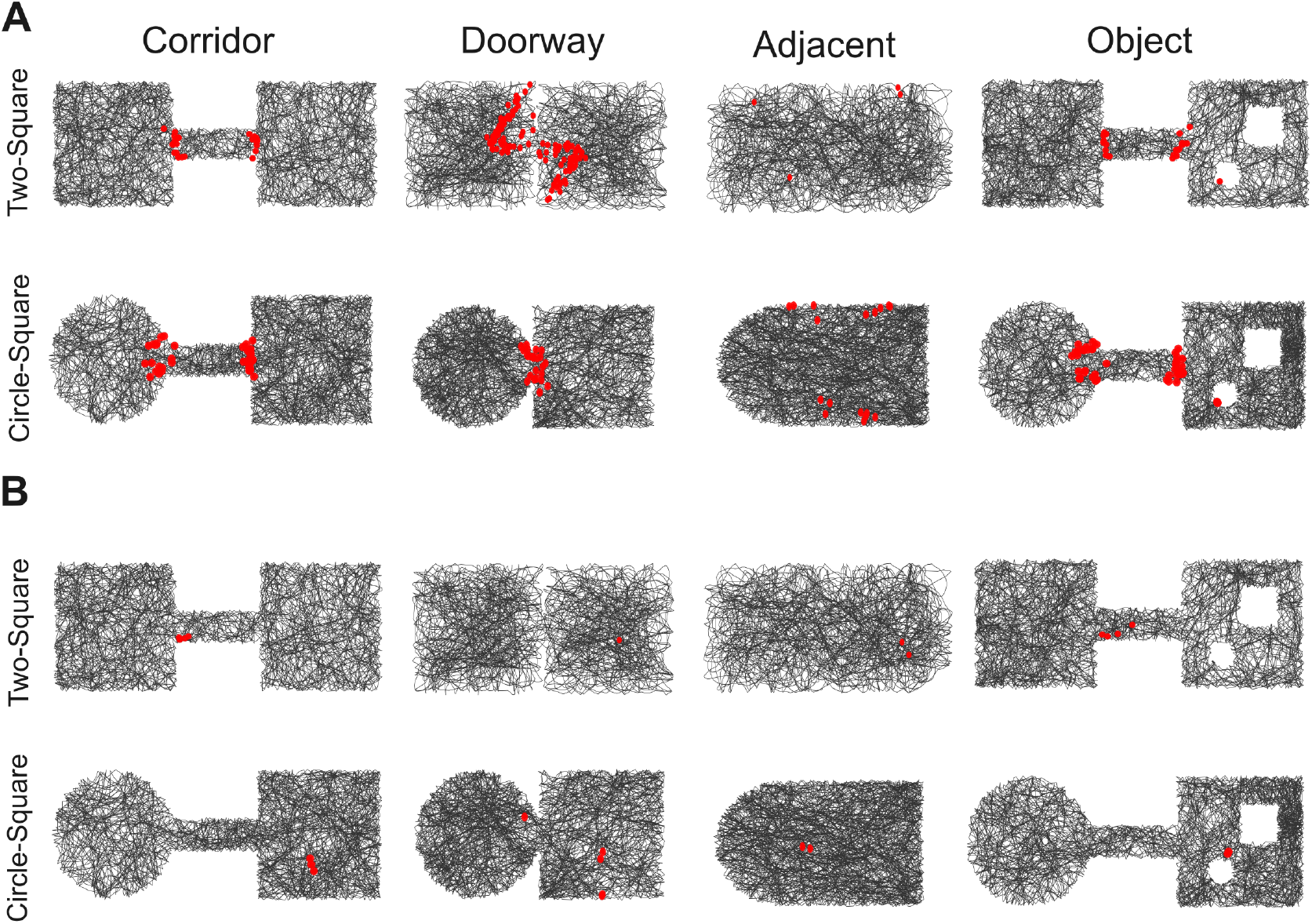
The influence of Disco axis cells on the event-boundary detection. The agent’s trajectory is shown in black. Red dots indicate time points at which the difference in discounted cumulative population activity (non-corner cells, corner cells, BVCs and axis cells) exceeds the threshold. The analysis was conducted on the proposed a population vector, M(*t*) (materials and methods). (**A-B**) Scaling factor of axis cells was set to *w*_*A*_ = 1 (A) and *w*_*A*_ = 100 (B). All other parameters were identical to those used in Fig. 6E.

**Fig. S20.**
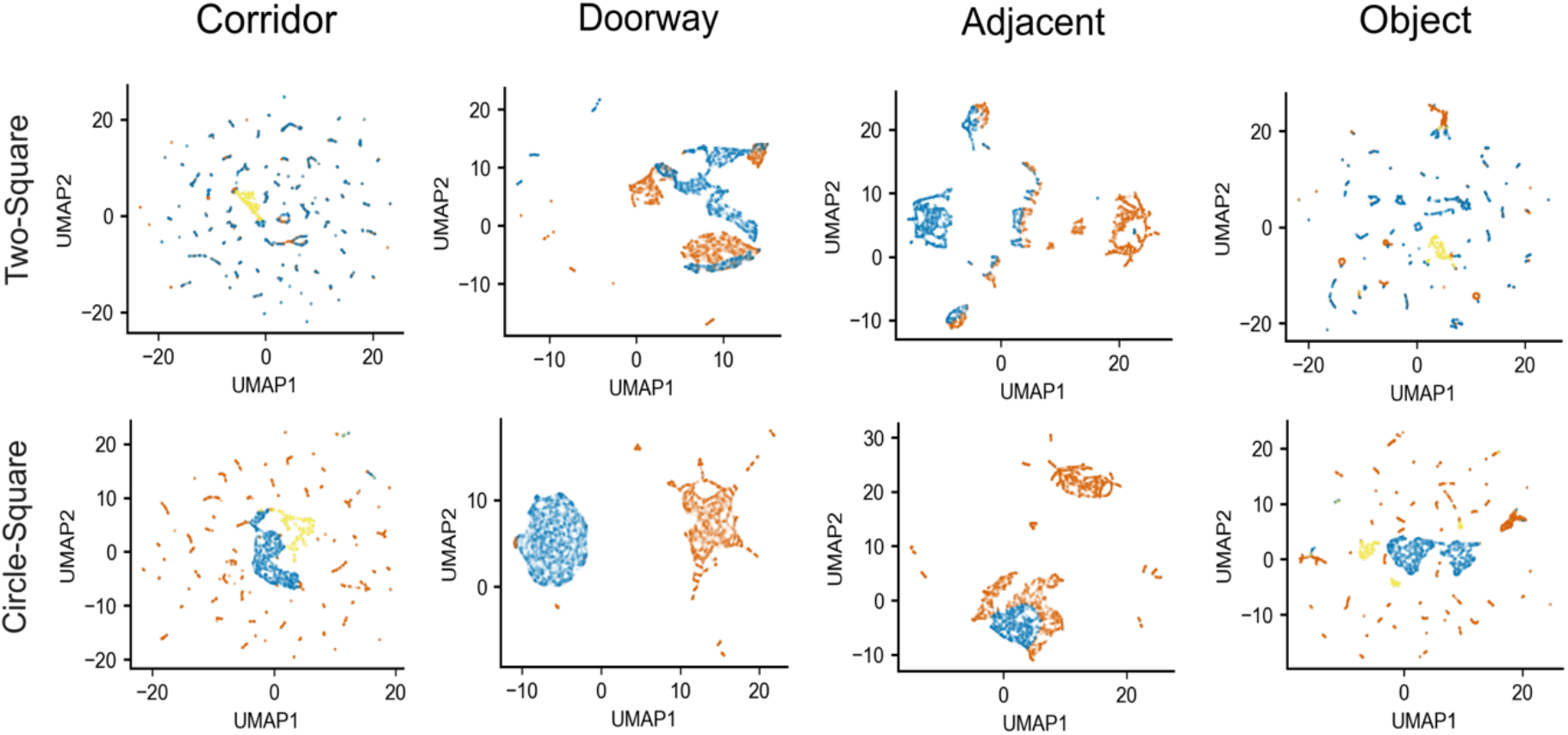
Additional dimension reduction method to analyze population activity. Additional dimensionality reduction analysis of population activity. Two-dimensional non-linear embeddings (UMAP) of population activity in the two-compartment arenas. Each dot represents the population state at a single spatial position, and arena regions corresponding to agent positions are color-coded as in Fig. 6.

**Fig. S21.**
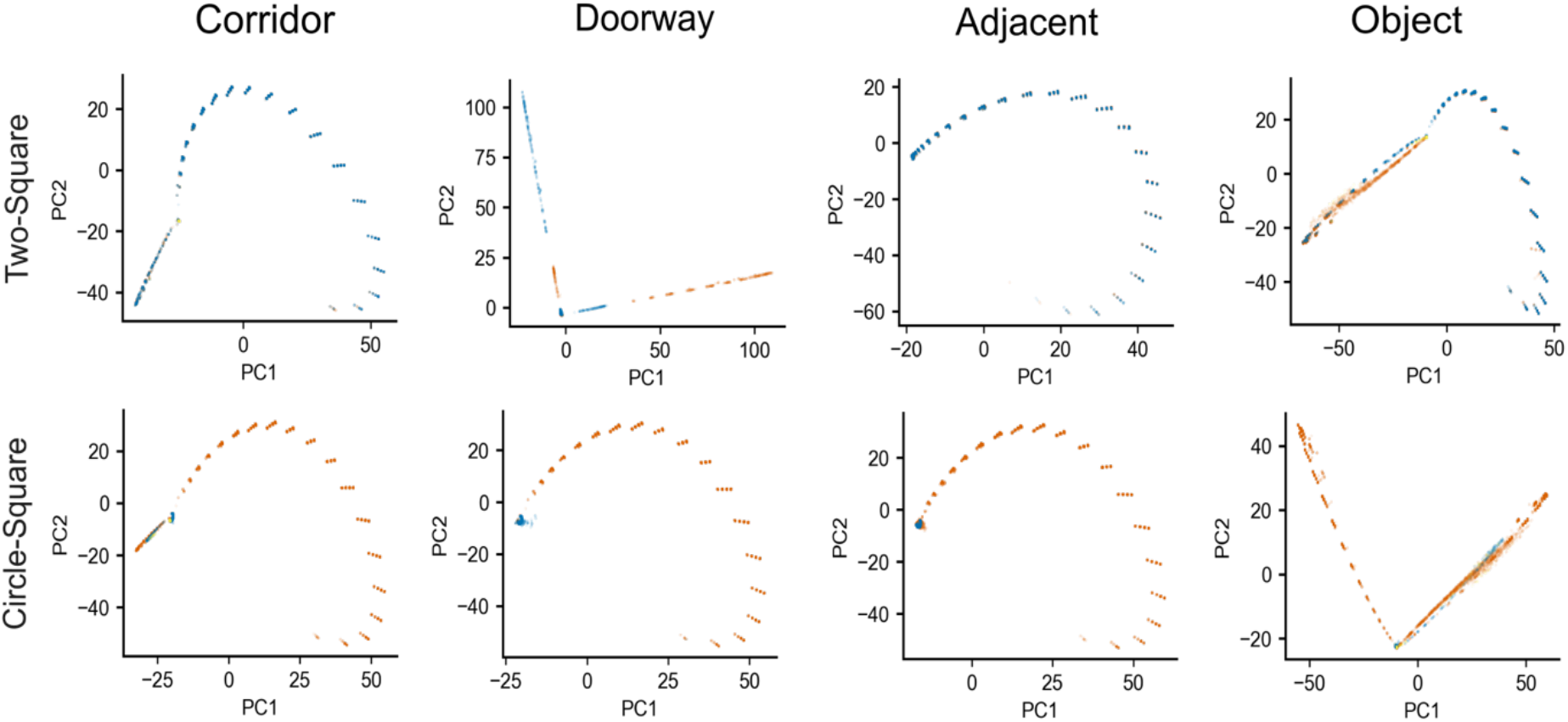
Additional dimension reduction method to analyze population activity. Two-dimensional linear embeddings (PCA) of population activity in the two-compartment arenas. Each dot represents the population state at a single spatial position, and arena regions corresponding to agent positions are color-coded as in Fig. 6.

Because we did not consider animal movement characteristics in open fields (Fig. 6A to G), Disco axis cells were not included in the analysis of the two-compartment simulations in the main text. Our simulated exploration trajectories exhibited a high degree of randomness, such that Disco axis cells did not exert a meaningful influence on the analysis results (fig. S18-19). We varied the scaling factor *w*_*A*_, which controls the relative contribution of Disco axis cells to population activity (Equation. (35)). When *w*_*A*_was small, Disco axis cells did not affect geometric shape encoding or event boundary detection. In contrast, large values of *w*_*A*_ led to randomized population representations and failed to detect the event boundary.

### E Simulation details

We used RatInABox (56) to simulate agent trajectories in multiple environments. The details of simulations involving Disco axis cells are as follows. For the open-field task, we simulated a circular arena with a radius of 30 cm for a duration of 30 min; For the linear task in the hairpin maze (dimensions shown in fig. S1), the agent traversed from the start point to the endpoint for 10 trials; For the triple ‘T’-track maze, we replicated the same environment configuration as the previous experiment (see details in Reference (19)) and simulated 10 trials for each of the eight linear exploration routes. All simulated one-dimensional linear trajectories were unidirectional, i.e., modeling well-trained animals proceeding from the task start point to the endpoint with no backtracking.

For simulations not involving Disco axis cell analyses, the details of the natural-environment simulations are provided in the main text (materials and methods; Fig. 4). The dimensions of all other open-field environments are shown in fig. S1, and the corresponding simulation durations are listed in table S2.

